# Deep-learning based identification, tracking, pose estimation, and behavior classification of interacting primates and mice in complex environments

**DOI:** 10.1101/2020.10.26.355115

**Authors:** Markus Marks, Jin Qiuhan, Oliver Sturman, Lukas von Ziegler, Sepp Kollmorgen, Wolfger von der Behrens, Valerio Mante, Johannes Bohacek, Mehmet Fatih Yanik

## Abstract

The quantification of behaviors of interest from video data is commonly used to study brain function, the effects of pharmacological interventions, and genetic alterations. Existing approaches lack the capability to analyze the behavior of groups of animals in complex environments. We present a novel deep learning architecture for classifying individual and social animal behavior, even in complex environments directly from raw video frames, while requiring no intervention after initial human supervision. Our behavioral classifier is embedded in a pipeline (SIPEC) that performs segmentation, identification, pose-estimation, and classification of complex behavior, outperforming the state of the art. SIPEC successfully recognizes multiple behaviors of freely moving individual mice as well as socially interacting non-human primates in 3D, using data only from simple mono-vision cameras in home-cage setups.

## Introduction

While the analysis of animal behavior is crucial for systems neuroscience^1^ and preclinical assessment of therapies, it remains a highly laborious and error-prone process. Over the last few years, there has been a surge in machine learning tools for behavioral analysis, including segmentation, identification, and pose estimation^2–11^. Although this has been an impressive feat for the field, a key element, the direct recognition of behavior itself, has been rarely addressed. Unsupervised analysis of behavior^12–17^ can be a powerful tool to capture the diversity of the underlying behavioral patterns, but the results of these methods do not align with human annotations and therefore require subsequent inspection^15^. There have been advances also in the supervised analysis of mouse behavior, using classifiers on top of pose-estimation generated features^18–21^ or manually defined features such as ellipses^22–25^. Sturman et. al.^20^ demonstrated that the classification of mouse behaviors using features generated from pose-estimation algorithms can outperform the behavioral classification performance of commercial systems. Yet, such pose-estimation-based behavior classification remains a labor-intensive and error-prone process as we show below. Moreover, pose estimation in primates is difficult to achieve with current methods^26^.

Here, we demonstrate a complementary approach for researchers who automatically seek to identify behaviors of interest. Our approach relies on the initial annotation of exemplar behaviors, i.e. snippets of video footage. These video snippets are subsequently used to train a Deep Neural Network (DNN) to subsequently recognize such particular behaviors in arbitrarily long videos and complex environments. To achieve this, we designed a novel DNN architecture, called SIPEC:BehaveNet, which uses raw videoframes as input and significantly outperforms a pose-estimation-based approach tested on a well-annotated mouse dataset and reaches human-level performances for counting grouped behavioral events. In addition to this behavioral classification network, we developed the first all-inclusive pipeline, called SIPEC, with modules for segmentation (SIPEC:SegNet), identification (SIPEC:IdNet), behavioral classification (SIPEC:BehaveNet), and pose estimation (SIPEC:PoseNet) of multiple and interacting animals in complex environments. This pipeline utilizes four DNNs operating directly on videos, developed and optimized for analyzing animal behavior and providing state-of-the-art performance. We use this pipeline to classify, for the first time, social interactions in home-caged primates from raw video frames and without needing to use any pose estimation.

SIPEC:SegNet is a Mask R-CNN architecture^27^, optimized to robustly segment animals despite occlusions, multiple scales, and rapid movement, and enables tracking of animal identities within a session. SIPEC:IdNet has a DenseNet^28^ backbone, that yields visual features, that are integrated over time through a gated-recurrent-unit network (GRU)^29,30^ to re-identify animals when temporal-continuity-based tracking does not work, for example when animals enter or exit a scene. This enables SIPEC to identify primates across weeks and to outperform the identification module of idtracker.ai^4^ both within-session and across sessions (see also Discussion) as well as primnet^31^. SIPEC:PoseNet performs top-down multi-animal pose estimation which we compared to DeepLabCut (DLC)^2^. SIPEC:BehaveNet uses an Xception^32^ network in combination with a temporal convolution network (TCN)^33,34^ to classify behavioral events directly from raw pixels. To rapidly train our modules, we use image augmentation^35^ as well as transfer-learning^36^, optimized specifically for each task. SIPEC enables researchers to identify behaviors of multiple animals in complex and changing environments over multiple days or weeks in 3D space, even from a single camera with relatively little labeling, in contrast to other approaches that use heavily equipped environments and large amounts of labelled data^8^.

To accelerate the reusability of SIPEC, we share the network weights among all four modules for mice and primates, which can be directly used for analyzing new animals in similar environments without further training or serve as pre-trained networks to accelerate training of networks in different environments.

## Results

Our algorithm performs segmentation (SIPEC:SegNet) followed by identification (SIPEC:IdNet), behavioral classification (SIPEC:BehaveNet) and finally pose estimation (SIPEC:PoseNet) from video frames (Fig. 1). These four artificial neural networks, trained for different purposes, can also be used individually or combined in different ways (Fig. 1a). To illustrate the utility of this feature, Fig. 1b shows the output of pipelining SIPEC:SegNet and SIPEC:IdNet to track the identity and location of 4 primates housed together (Fig. 1b, Supp. Video 1). Fig. 1c shows the output of pipelining SIPEC:SegNet and SIPEC:PoseNet to do multi-animal pose estimation in a group of 4 mice.

**Fig. 1.**
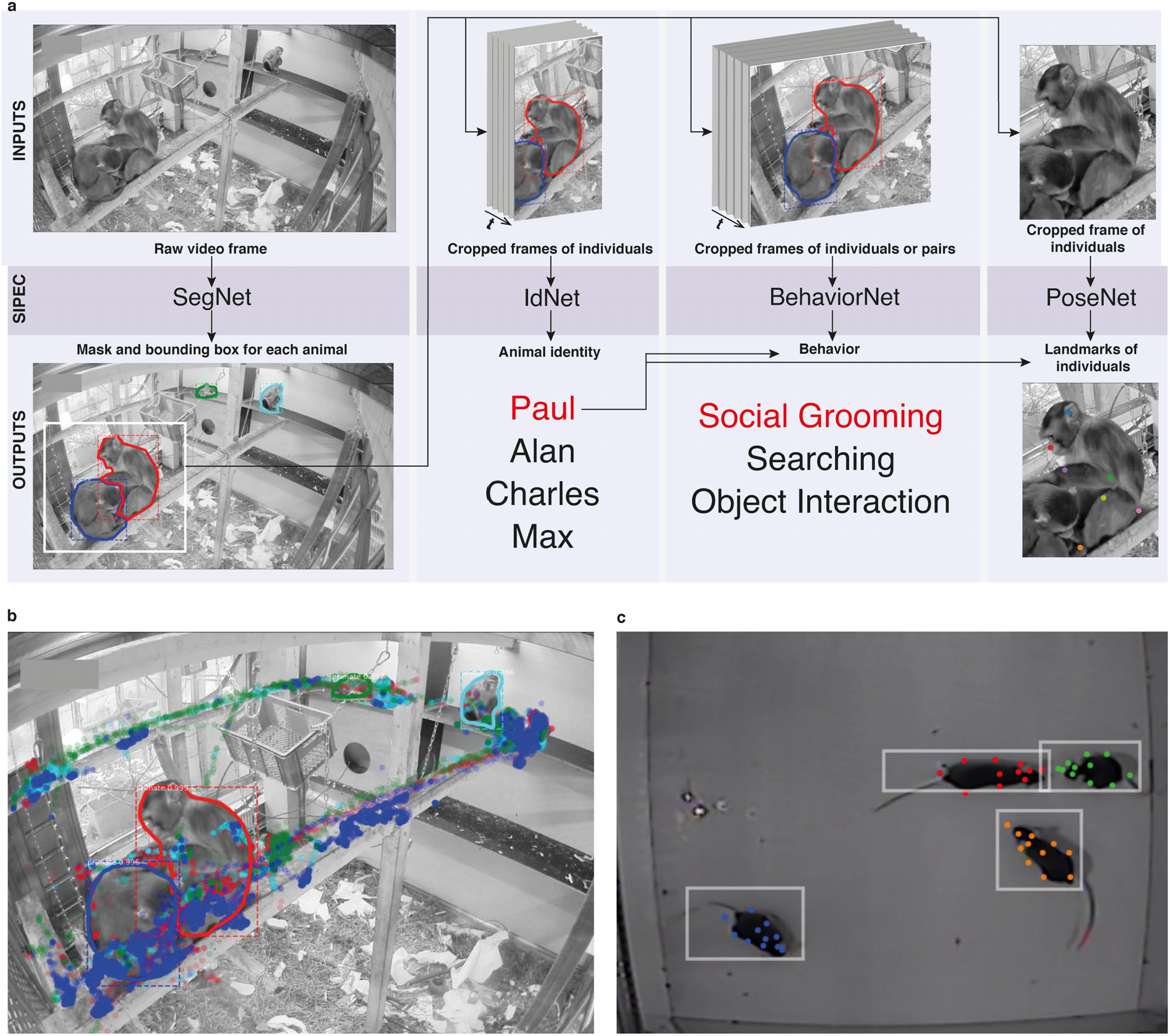
Overview of the SIPEC workflow and modules. a) From a given video, instances of animals are segmented with the segmentation network (SIPEC:SegNet), indicated by masked outline as well as bounding boxes. Subsequently, individuals are identified using the identification network (SIPEC:IdNet). For each individual, the pose and behavior can be estimated/classified using the pose estimation network (SIPEC:PoseNet) and the behavioral identification network (SIPEC:BehaveNet), respectively. b) Outcome of SIPEC:SegNet, and SIPEC:IdNet modules are overlaid on a representative videoframe. Time-lapsed positions of individual primates (center of mass) are plotted as circles with respective colors. c) Outputs of SIPEC:SegNet (boxes) and SIPEC:PoseNet (colored dots) on a representative videoframe of mouse open-field data.

### Segmentation module SIPEC:SegNet

SIPEC:SegNet (see Methods, Supp. Fig. 12) is based on the Mask-RCNN architecture^27^, which we optimized for analyzing multiple animals and integrated into SIPEC. We further applied transfer learning^36^ onto the weights of the Mask-RCNN ResNet-backbone^37^ pre-trained on the Microsoft Common Objects in Context (COCO dataset)^38^ (see Methods for SIPEC:SegNet architecture and training). Moreover, we applied image augmentation^35^ to increase network robustness against invariances, e.g. rotational invariance and therefore increase generalizability.

#### Segmentation performance on individual mice and groups of 4

We first examined the performance of SIPEC:SegNet on top-view video recordings of individual mice, behaving in an open-field test (OFT). While segmenting black mice on a blank background could be achieved by thresholding alone, we still included this task for completeness. 8 mice were freely behaving for 10 minutes in the TSE Multi Conditioning System’s OFT arena, previously described in Sturman et al.^20^. We labeled the outlines of mice in a total of 23 frames using the VGG image annotator^39^ from videos of randomly selected mice. To evaluate the performance, we used 5-fold cross-validation (CV). We assessed the segmentation performance on images of individual mice, where SIPEC:SegNet achieved a mean-Average Precision (mAP) of 1.0 ± 0 (mean ± s.e.m., see Methods for metric details). We performed a videoframe ablation study to determine how many labeled frames (outline of the animal, see Supp. Fig. 1) are needed for SIPEC:SegNet to reach peak performance (Supp. Fig. 2). While randomly selecting an increasing amount of training frames, we measured performance using CV. For single-mouse videos, we find that our model achieves 95% of mean peak performance (mAP of 0.95 ± 0.05) using as few as a total of 3 labeled frames for training. To the existing 23 labeled single-mouse frames, we added 57 labeled 4-plex frames, adding to a total of 80 labeled frames. Evaluated on a 5-fold CV, SIPEC:SegNet achieves an mAP of 0.97 ± 0.03 (Fig. 2b). For segmentation in groups of 4 mice, we performed an ablation study as well and found that SIPEC:SegNet achieves better than 95% of the mean peak performance (mAP of 0.94 ± 0.05) using as few as only 16 labeled frames. To assess the overlap between prediction and ground truth, we report IoU and dice coefficient metrics as well (Fig. 2b).

**Fig. 2.**
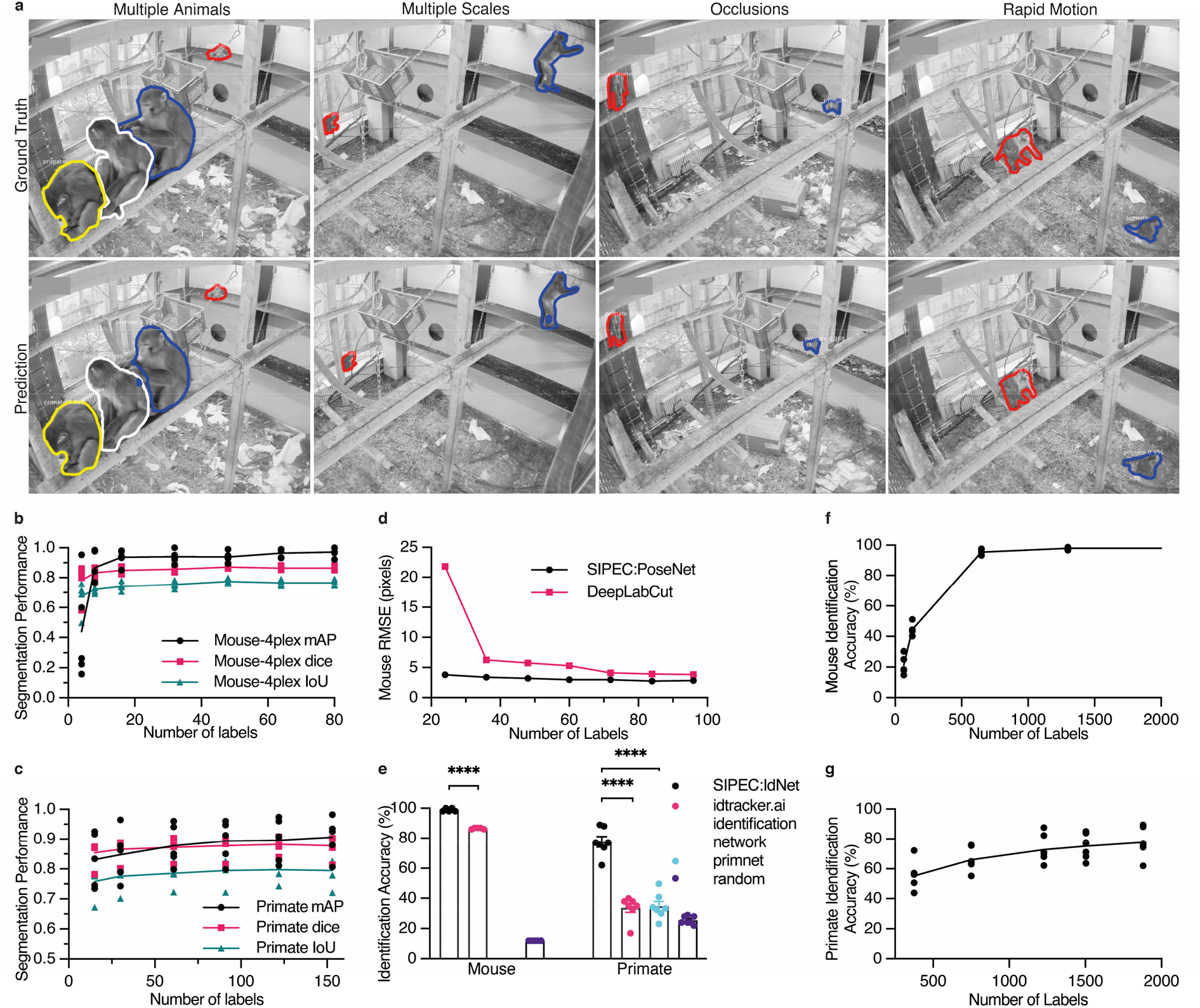
Performance of the segmentation (SIPEC:SegNet), pose estimation (SIPEC:PoseNet), and identification (SIPEC:IdNet) modules under demanding video conditions and using few labels. a) Qualitative comparison of ground truth (top row) versus predicted segmentation masks (bottom row) under challenging conditions; multiple animals, at varying distances from the camera, under strong visual occlusions, and in rapid motions. b) For mice, SIPEC:SegNet performance in mAP (mean average precision), dice (dice coefficient), and IoU (intersection over union) as a function of the number of labels. The lines indicate the means for 5-fold CV while circles, squares, triangles indicate the mAP, dice, and IoU, respectively, for individual folds. c) For primates, SIPEC:SegNet performance in mAP, dice, and IoU as a function of the number of labels. The lines indicate the means for 5-fold CV while circles, squares, triangles indicate the mAP, dice, and IoU, respectively, for individual folds. d) The performance of SIPEC:PoseNet in comparison to DeepLabCut measured as RMSE in pixels on single mouse pose estimation data. e). Comparison of identification accuracy for SIPEC:IdNet module, idtracker.ai^4^, primnet^31^ and randomly shuffled labels (chance performance). 8 videos from 8 individual mice and 7 videos across 4 different days from 4 group-housed primates are used. f) For mice, the accuracy of SIPEC:IdNet as a function of the number of training labels used. The black lines indicate the mean for 5-fold CV with individual folds displayed. g) For primates, the accuracy of SIPEC:IdNet as a function of the number of training labels used. The black lines indicate the mean for 5-fold CV with individual folds displayed. All data is represented by mean, showing all points.

#### Segmentation performance of groups of primates

To test SIPEC:SegNet for detecting instances of primates within a group, we annotated 191 frames from videos on different days (Day 1, Day 9, Day 16, Day 18). As exemplified in Fig. 2a, the network handles even difficult scenarios very well: representative illustrations include ground-truth as well as predictions of moments in which multiple primates are moving rapidly while strongly occluded at varying distances from the camera. SIPEC:SegNet achieved a mAP of 0.91 ± 0.03 (mean ± s.e.m.) using 5-fold CV. When we performed the previously described ablation study, SIPEC:SegNet achieved 95% of mean peak performance (mAP of 0.87 ± 0.03) with only 30 labeled frames (Fig. 2b). To assess the overlap between prediction and ground truth, we report IoU and dice coefficient metrics as well (Fig. 2c).

### Pose estimation module SIPEC:PoseNet

We also added a pose estimation network, built on an encoder-decoder architecture^40^ with an EfficientNet^41^ backbone, to SIPEC for performing pose estimation (SIPEC:PoseNet) (see Methods, Supp. Fig. 11). SIPEC:PoseNet can be used to perform pose estimation on *N* animals (with *N* the total number of animals or less), yielding *K* different coordinates for previously defined landmarks on each animal’s body. The main advantage of SIPEC:PoseNet in comparison to previous approaches is that it receives its inputs from SIPEC:SegNet (top-down pose estimation): While bottom-up approaches such as DLC^2^ require grouping of pose estimates to individuals, our top-down approach makes the assignment of pose estimates to individual animals trivial, as inference is performed on the masked image of an individual animal and pose estimates within that mask are assigned to that particular individual (Fig. 1c). We labeled frames with 13 standardized body parts of individual mice in an OFT similarly to Sturman et. al.^20^ to train and test the performance of SIPEC:PoseNet against that of DLC^2^. SIPEC:PoseNet achieves a Root-Mean-Squared-Error (RMSE) (see Methods) of 2.9 pixels in mice (Fig. 2d) for a total of 96 labeled training frames, while DLC^2^ achieves a 3.9 pixel RMSE^2^. Previously published pose estimation methods for single animals can easily be substituted into our pipeline to perform multi-animal pose estimation in conjunction with SIPEC:SegNet.

### Identification module SIPEC:IdNet

The identification network (SIPEC:IdNet) (see Methods, Supp. Fig. 10) allows the determination of the identity of individual animals. Given SIPEC:IdNet receives input as a series (*T* time points) of cropped images of *N* (with *N* the total number of animals or less) individuals from SIPEC:SegNet, the output of SIPEC:IdNet are *N* identities. The input images from SIPEC:SegNet are scaled to the same average size (see Methods) before being fed into SIPEC:IdNet. We designed a feedforward classification neural network, which utilizes a DenseNet^28^-backbone pre-trained on ImageNet^42^. This network serves as a feature-recognition network on single frames. We then utilize past and future frames by dilating the mask around the animal with each timestep. The outputs of the feature-recognition network on these frames are then integrated over *T* timesteps using a GRU (see Methods for architecture and training details). SIPEC:IdNet can integrate information from none to many temporally-neighboring frames based on a particular application’s accuracy and speed requirements. We used spatial area dropout augmentations to increase robustness against occlusions^43^. We developed an annotation tool for a human to assign identities of individual animals, in a multi-animal context, to segmentation masks in videoframes, which capture primates from different perspectives (Supp. Fig. 3). This tool was used for annotating identification data in the following sections. Below we compared SIPEC:IdNet’s performance to that of the current state-of-the-art i.e. the identification module of idTracker.ai^4^ and the primnet^31^ network for primate re-identification. primnet^31^ relies on faces of individuals being clearly visible for re-identification, which in our case is not possibe for most of the video frames. idTracker.ai^4^ is a self-supervised algorithm for tracking the identity of individual animals within a single session. Particularly in complex or enriched home-cage environments, where animals are frequently obstructed as they move underneath/behind objects or enter/exit the scene and background or lighting conditions change constantly, temporally based tracking and identification as idtracker.ai performs it becomes impossible. We evaluated the identification performance of SIPEC:IdNet across sessions with the identification module of idTracker.ai, providing each network with identical training and testing data. While idtracker.ai behaves self-supervised, the identification module it uses to distinguish animals is trained with the labels generated by idTracker.ai’s cascade algorithm in a supervised fashion. Apart from re-identifying animals across sessions using SIPEC:IdNet, SIPEC:SegNet segmentation masks can be used via greedy-mask matching (see Methods) to track the identities of animals temporally as well (Supp. Videos 2-4) or to smooth the outputs of SIPEC:IdNet as a secondary step, that can boost performance for continuous video sequences, but this advantage was not used in the following evaluations for mice and primates.

#### Identification of mice in an open-field test

We first evaluated the performance of SIPEC:IdNet in identifying 8 individual mice. We acquired 10-minute-long videos of these mice behaving in the previously mentioned OFT (see Methods for details). While for the human observer, these mice are difficult to distinguish (Supp. Fig. 4), our network copes rather well. We used 5-fold CV to evaluate the performance, i.e. splitting the 10-minute videos into 2-minute long ones, while using one fold for testing and the rest to train the network Since this data is balanced, we use the accuracy metric for evaluation. We find that SIPEC:IdNet achieves 99 ± 0.5 % (mean and s.e.m.) accuracy, while the current state of the art idTracker.ai^4^ only achieves 87 ± 0.2 % accuracy (Fig. 2e). The ablation study shows that only 650 labeled frames (frame and identity of the animal) are sufficient for the SIPEC:IdNet to achieve 95% of its mean peak performance (Fig. 2f). We tested how this performance translates to identifying the same animals during the subsequent days (Supp. Fig. 5). We find that identification performance is similarly high on the second day 86 ± 2 %, using the network trained on day 1. Subsequently, we tested identification robustness with respect to the interventions on day 3. Following a forced swim test, the identification performance of SIPEC:IdNet, trained on data of day 1, dropped dramatically to 4 ± 2 %. This indicates that features utilized by the network to identify the mice are not robust to this type of intervention, i.e. their behavior and outlook is altered by the stress and residual water on the fur significantly.

#### Identification of individual primates in a group

To evaluate SIPEC: IdNet’s performance on identifying individual primates within a group, we used the SIPEC:SegNet-processed videos of the 4 macaques (see Section “Segmentation performance of groups of primates”). We annotated frames from 7 videos taken on different days, with each frame containing multiple individuals, yielding approximately 2200 labels for cutouts of individual primates. We used leave-one-out CV with respect to the videos in order to test SIPEC:IdNet generalization across days. Across sessions SIPEC:IdNet reaches an accuracy of 78 ± 3 % (mean ± s.e.m.) while idTracker.ai^4^ achieves only 33 ± 3 % and primnet^31^ 34 ± 3 % (Fig. 2e), where the human expert (i.e. ground truth) had the advantage of seeing all the video frames and the entire cage (i.e. the rest of the primates). We did a separate evaluation of the identification performance on “typical frames” i.e., the human expert can correctly identify the primates using single frames. In this case, SIPEC:IdNet achieved a performance of 86 ± 3 (Supp. Fig. 6). The identification labels can then be further enhanced by greedy mask-match-based tracking (see Methods for details). Supp. Video 1 illustrates the resulting performance on a representative video snippet. We perform here an ablation study as well, which yields 95% of mean peak performance at 1504 annotated training samples (Fig. 2g).

### Behavioral classification module SIPEC:BehaveNet

SIPEC:BehaveNet (see Methods, Supp. Fig. 13) offers researchers a powerful means to recognize specific animal behaviors directly from raw pixels using a single neuronal net framework. SIPEC:BehaveNet uses video frames of *N* individuals over *T* time steps to classify the animals’ actions. The video frames of the *N* individuals are generated by SIPEC:SegNet. If only a single animal is present in the video, SIPEC:BehaveNet can be used directly without SIPEC:SegNet. We use a recognition network to extract features from single frames analysis, based on the Xception^32^ network architecture. We initialize parts of the network with ImageNet^4^ weights. These features are then integrated over time by a TCN^33,34^ to classify the animal’s behavior in each frame (see Methods for architecture and training details).

#### SIPEC behavior recognition outperforms DLC-based approach

We compare our raw-pixel-based approach to Sturman et al.^20^, who recently demonstrated that they can classify behavior based on DLC^2^ generated features. On top of a higher classification performance with fewer labels, SIPEC:BehaveNet does not require annotation and training for pose estimation if the researcher is interested in behavioral classification alone. The increased performance with fewer labels comes at the cost of a higher computational demand since we increased the dimensionality of the input data by several orders of magnitude (12 pose estimates vs. 16384 pixels). We used the data and labels from Sturman et al.^20^ on 20 freely behaving mice in an OFT to test our performance. The behavior of these mice was independently annotated by 3 different researchers on a frame-by-frame basis using the VGG video annotation tool^39^. Annotations included the following behaviors: supported rears, unsupported rears, grooming and none (unlabeled/default class). While Sturman et al.^20^ evaluated the performance of their behavioral event detection by averaging across chunks of time, evaluating the frame-by-frame performance is more suitable for testing the actual network performance since it was trained the same way. Doing such frame-by-frame analysis shows that SIPEC:BehaveNet has fewer false positives as well as false negatives with respect to the DLC-based approach of Sturman et al. ^20^. We illustrate a representative example of the performance of both approaches for each of the behaviors with their respective ground truths (Fig. 3a). We further resolved spatially the events that were misclassified by Sturman et al., that were correctly classified by SIPEC:BehaveNet and vice versa (Fig. 3b). We calculated the percentage of mismatches, that occurred in the center or the surrounding area. For grooming events mismatches of Sturman et al.^20^ and SIPEC:BehaveNet occurs similarly often in the center 41 ± 12 % (mean and s.e.m.) and 42 ± 12 % respectively. For supported and unsupported rearing events Sturman et al.^20^ has more mismatches occurring in the center compared to SIPEC:BehaveNet (supported rears: 40 ± 4 % and 37 ± 6 %, unsupported rears: 12 ± 2 % and 7 ± 2 %). This indicates that the misclassifications of the pose estimation-based approach are more biased towards the center than the ones of SIPEC:BehavNet. To quantify the behavioral classification over the whole time course of all videos of 20 mice, we used leave-one-out CV (Fig. 3c). We used macro-averaged F1-score as a common metric to evaluate a multi-class classification task and Pearson correlation (see Methods for metrics) to indicate the linear relationship between the ground truth and the estimate over time. For the unsupported rears/grooming/supported rears behaviors SIPEC:BehaveNet achieves F1-Scores of 0.6 ± 0.16/0.49 ± 0.21/0.84 ± 0.04 (values reported as mean ± s.e.m.) respectively, while the performance of the manually intensive Sturman et al.^20^’s approach reaches only 0.49 ± 0.11/0.37 ± 0.2/0.84 ± 0.03, leading to a significantly higher performance of SIPEC:BehaveNet for the unsupported rearing (F1: p=1.689×10^−7^, Wilcoxon paired-test was used as recommended^44^) as well as the grooming (F1: p=6.226×10^−4^) behaviors. While we see a higher precision only in the classification of supported rears in the DLC-based approach, SIPEC:BehaveNet has an improved recall for the supported rears as well as improved precision and recall for the other behaviors (Supp. Fig. 7a). As expected, more stereotyped behaviors with many labels like supported rears yield higher F1. In comparison, less stereotypical behaviors like grooming with fewer labels have lower F1 for SIPEC:BehaveNet and the DLC-based approach. Additionally, we computed the mentioned metrics on a dataset with shuffled labels to indicate chance performance for each metric as well as computed each metric when tested across human annotators to indicate an upper limit for frame-by-frame behavioral classification performance (Supp. Fig. 7b). While the overall human-to-human F1 is 0.79 ± 0.07 (mean ± s.e.m.), SIPEC:BehaveNet classifies with an F1 of 0.71 ± 0.07. We then grouped behaviors by integrating the classification over multiple frames as described in Sturman et al.^20^. This analysis results in a behavior count per video. For these per video behavior counts, we found no significant difference between human annotators, SIPEC:BehaviorNet and Sturman et al.^20^ (Tukey’s multiple comparison test, Supp. Fig. 15). Such classification and counting of specific behaviors per video are commonly used to compare the number of occurrences of behaviors across experimental groups. Using such analysis, Sturman et al.^20^ demonstrate how video-based analysis outperforms commonly used commercial systems. Moreover, we also tested combining the outputs of pose estimation-based classification together with the raw-pixel model (Combined Model in Methods, Supp. Fig. 7). Lastly, we performed a frame ablation study and showed that SIPEC:BehaveNet needs only 114 minutes, less than 2 hours, of labeled data to reach peak performance in behavior classification (Fig. 3d).

**Fig. 3.**
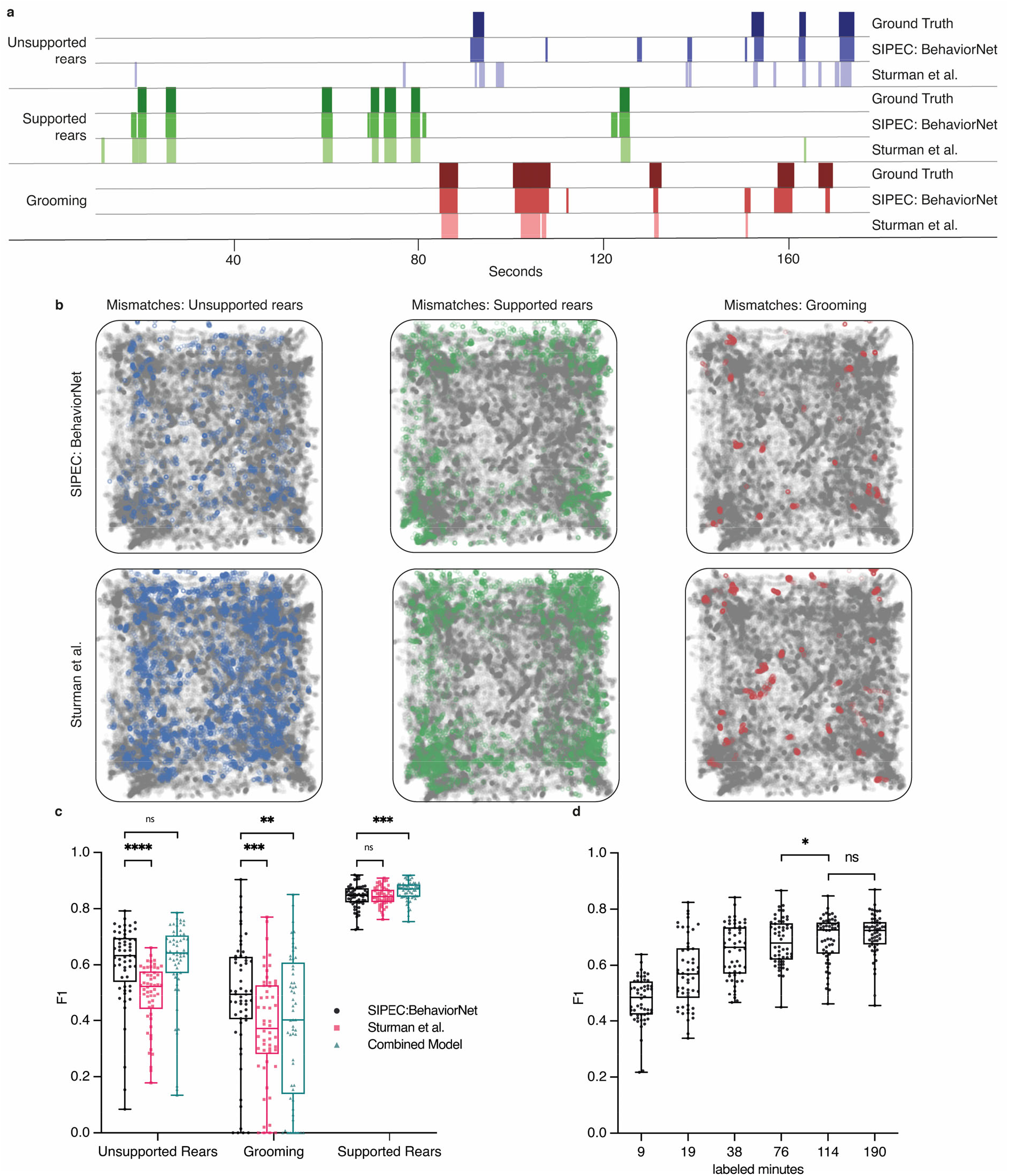
SIPEC:BehaveNet outperforms pose-estimation (DeepLabCut) based approach (Sturman et al.^20^). a) Comparison of behavioral classification by human annotator (ground truth), SIPEC:BehaveNet, and Sturman et al.^20^ b) Errors in the classification of mouse behavior in the open arena for SIPEC:BehaveNet versus Sturman et al. Each colored dot represents a behavioral event that is incorrectly classified by that method (while correctly classified by the other) with respect to the ground truth. none-classified (background class) positions of mice are indicated as grey dots. c) Frame-by-frame classification performance per video (n=20 mice) compared to ground truth. d) SIPEC:BehaveNet classification performance as a function of labeled minutes. All data is represented by a Tukey box-and-whisker plot, showing all points. Wilcoxon paired test:* p <= 0.05; *** p <= 0.001; **** p <= 0.0001.

### Socially interacting primate behavior classification

We used the combined outputs of SIPEC:SegNet and SIPEC:IdNet, smoothed by greedy match-based tracking, to generate videos of individual primates over time (see Methods for details). To detect social events, we used SIPEC:SegNet to generate additional video events covering “***pairs***” of primates. An interaction event was detected whenever the masks of individual primates came sufficiently close (see Methods). We were able to rapidly annotate these videos again using the VGG video annotation tool^39^ (overall 80 minutes of video are annotated from 3 videos, including the individual behaviors of object interaction, searching, **social grooming** and none (background class)). We then trained SIPEC:BehaveNet to classify individuals’ frames and merged frames of pairs of primates socially interacting over time. We used grouped 5-fold stratified CV over all annotated video frames, with labeled videos being the groups. Overall SIPEC:BehaveNet achieved a macro-F1 of 0.72 ± 0.07 (mean ± s.e.m.) across all behaviors (Fig. 4a). This performance is similar to the earlier mentioned mouse behavioral classification performance. The increased variance compared to the classification of mouse behavior is expected as imaging conditions, as previously mentioned, are much more challenging and primate behaviors are much less stereotyped compared to mouse behaviors. This can be likely compensated with more training data.

**Fig. 4.**
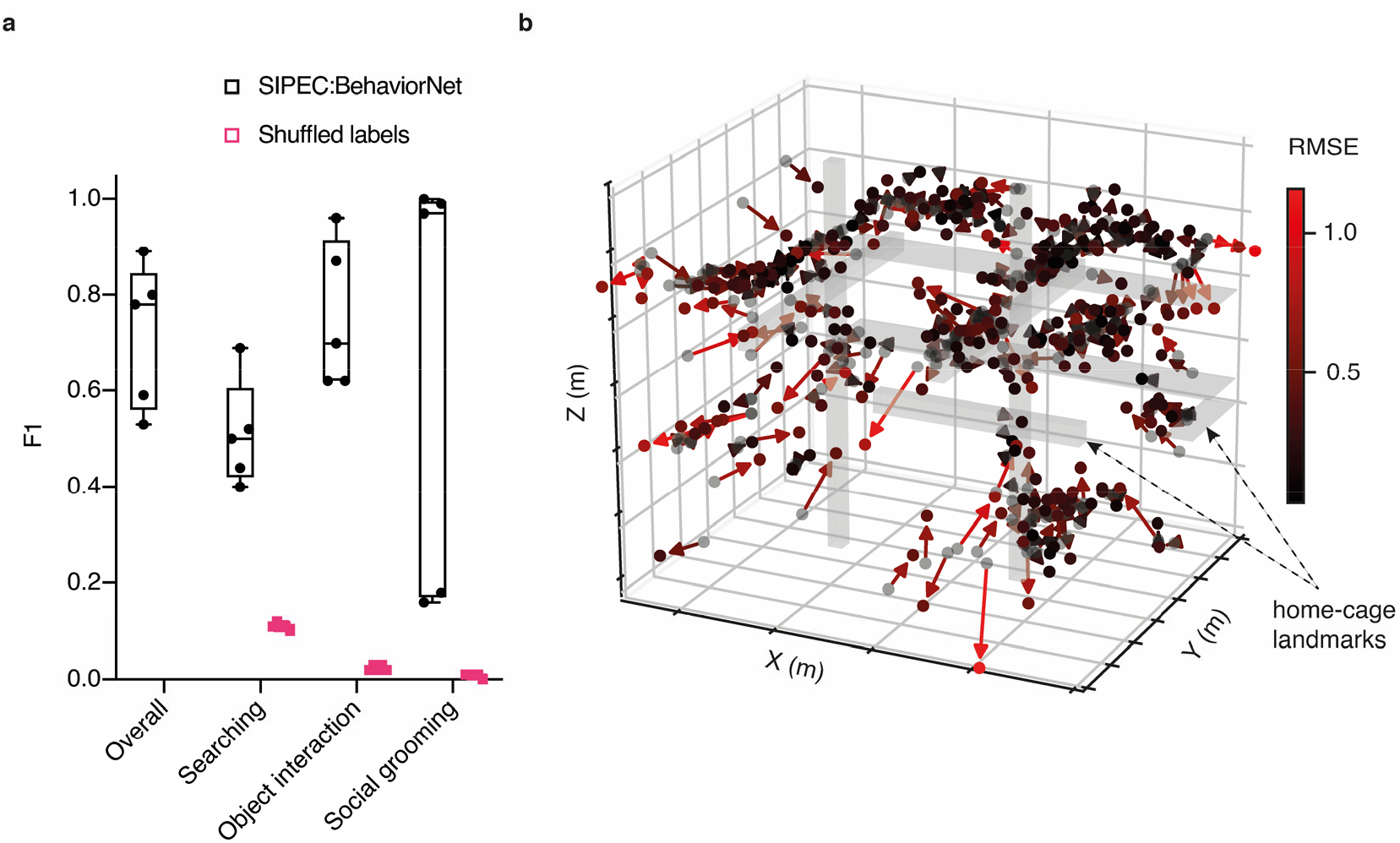
SIPEC can recognize social interactions of multiple primates and infer their 3D positions using a single camera. a) Performance of SIPEC:BehaveNet for individual and social behaviors with respect to ground truth evaluated using grouped 5-fold CV. Behaviors include searching, object interaction and social grooming; while the performance is measured using F1. F1 on shuffled labels is included for comparison. All data is represented by a minimum-to-maximum box-and-whisker plot, showing all points. b) Evaluation of 3D position estimates of primates in home-cage. Black spots mark annotated positions (n=300) while predicted positions are marked as red-hued spots at the end of the solid arrows (color-coded using a red gradient with brighter red indicating higher RMSE of predicted to true position).

### Tracking position of primates in 3D without stereo-vision

By performing SIPEC:SegNet and SIPEC:IdNet inference on a full one-hour video, we built a density map of positions of individuals within the husbandry (Fig. 1a). Without stereo-vision, one cannot optically acquire depth information. Instead, we used the output masks of SIPEC:SegNet and annotated the positions of the primates in 300 frames using a 3D model (Supp. Fig. 8). Subsequently, we generated 6 features using Isomap^45^ and trained a multivariate linear regression model to predict the 3D positions of the primates (Fig. 4b). Using 10-fold CV, our predicted positions using only single camera have an overall RMSE of only 0.43 ± 0.01 m (mean ± s.e.m.), that is of 0.27 ± 0.01 m in x-direction or 6% error w.r.t the room dimension in x-direction; 0.26 ± 0.01 m / 7% and 0.21 ± 0.01 m / 7% for the y and z coordinates respectively. If an annotation is impossible, quasi depth estimates can be calculated through the mask size alone and correlate highly with the actual depth (Supp. Fig. 14).

## Discussion

We have presented SIPEC, a novel pipeline, using specialized deep neural networks to perform segmentation, identification, behavioral classification, and pose estimation on individual and interacting animals. With SIPEC we address multiple key challenges in the domain of behavioral analysis. Our **SIPEC:SegNet** enables the segmentation of animals with only 3-30 labels (Fig. 2a,b,c). In combination with greedy-mask matching, SIPEC:SegNet can be used to track animals’ identities within one session similar to idtracker.ai, but even in complex environments with changing lighting conditions, where idtracker.ai fails (Supp. Video 1).

Subsequently, **SIPEC:BehaveNet** enables animal behavior recognition directly from raw video data. Raw-video classification has the advantage of not requiring pre-processing adjustments or feature engineering to specific video conditions. Moreover, we show that learning task-relevant features directly from the raw video can lead to better results than pose-estimation-based approaches which train a classifier on top of the detected landmarks. In particular, we demonstrate that our network outperforms a state-of-the-art pose estimation approach^13^ on a well-annotated mouse behavioral dataset (Fig. 3) and reaches human-level performance for counting behavioral events (Supp. Fig. 15). Thus, pose-estimation can be skipped if researchers are solely interested in classifying behavior. We note that our raw-pixel approach increases the input-dimensionality of the behavior classification network and therefore uses more computational resources and is slower than pose-estimation-based approaches.

**SIPEC:IdNet** identifies primates in complex environments across days with high accuracy. SIPEC:SegNet enhances SIPEC:IdNet’s high identification performance through mask-matching-based tracking and integration of identities through time. We demonstrate that identification accuracy is significantly higher than that of the identification module of state-of-art idtracker.ai and primnet^31^ (Fig. 2e). We note, however, that identification using deep nets is not robust to interventions that affect mice’s appearance strongly immediately after the intervention (such as forced swim test, Supp. Fig. 5). However, even without any interventions, expert human observers have difficulty identifying mice of such similar size and color. The effects of different interventions on the recognition performances of deep net architectures should be studied in the future. Finally, **SIPEC:PosNet** enables top-down pose estimation of multiple animals in complex environments, making it easy to assign pose estimates to individual animals with higher performance than DLC (Fig. 2d).

All approaches are optimized through augmentation and transfer learning, significantly speeding up learning and reducing labeling compared to the other approaches we tested on the mouse and non-human primate datasets. We also performed ablation studies for each of the networks to estimate the number of labels necessary for successful training. The number of labels necessary can change depending on the dataset, for example, if the background, etc. are more complex each network could require more annotated frames to be trained successfully. To perform well under the complex video conditions for non-human primates, SIPEC:SegNet needs about 30 labels, SIPEC:IdNet about 1500 labels and SIPEC:BehaveNet less than 2 hours of annotated video (Fig. 2c,g; Fig. 4a).

SIPEC can be used to study the behavior of primates and their social interactions over longer periods in a naturalistic environment, as we demonstrated for social grooming (Fig. 4a). In addition, after initial training of SIPEC modules, they can automatically output a behavioral profile for each individual in a group, over days or weeks and therefore also be used to quantify the changes in behaviors of individuals in social contexts over time. Since SIPEC is fully supervised, it may be difficult to scale it to large colonies with hundreds of animals, such as bees and ants. However, SIPEC is well suited for most other animal species beyond insects.

Finally, we show how SIPEC enables 3D localization and tracking from a single-camera view, yielding an off-the-shelf solution for home-cage monitoring of primates, without the need for setting stereo-vision setups (Fig. 4b). Estimating the 3D position requires the experimenter to create a 3D model and annotate 3D data. However, we show a quasi-3D estimate can be generated directly from the mask size, without manual annotation, that correlates highly with the actual position of the animal (Supp. Fig. 14).

Behaviors which were not recognized and annotated by the researcher and therefore not learned by the neural network could be picked up using complementary unsupervised approaches^12,13^. The features-vectors, embedding individual behaviors, created by SIPEC:BehaveNet can be used as input to unsupervised approaches, which can help align the outputs of unsupervised approaches with human annotation. Moreover, the output of other modules (SIPEC:SegNet, SIPEC:IdNet and SIPEC:PoseNet) can also be used after such unsupervised approaches to analyse individual animals.

## Data Availability

Mouse data from Sturman et al.^20^ is available under https://zenodo.org/record/3608658. Primate data is available upon reasonable request from authors. Exemplary data for training is available through our github repository.

## Code Availability

We provide the code for SIPEC at https://github.com/SIPEC-Animal-Data-Analysis/SIPEC (https://doi.org/10.5281/zenodo.5927367) and the GUI for the identification of animals https://github.com/SIPEC-Animal-Data-Analysis/idtracking_gui.

## Acknowledgments

This project was funded by the Swiss Federal Institute of Technology (ETH) Zurich and the European Research Council (ERC) under the European Union’s Horizon 2020 research and innovation program (grant agreement No 818179), SNSF (CRSII5_198739/1 to MFY; 310030_172889/1 to JB, PP00P3_157539 to VM) ETH Research Grant (ETH-20 19-1 to JB), 3RCC (OC-2019-009 to JB and MFY), the Simons Foundation (awards 328189 and 543013 to VM) and the Botnar Foundation (to JB). We would like to thank Petra Tornmalm and Victoria de La Rochefoucauld for annotating primate data and feedback on primate behavior. We would like to thank Paul Johnson, Baran Yasar, Bifeng Wu, and Aagam Shah for helpful discussions and feedback.

## Author contributions

M.M. developed, implemented, and evaluated the SIPEC modules and framework. J.Q. developed segmentation filtering, tracking, and 3D-estimation. M.M., W.B., and M.F.Y. wrote the manuscript. M.M., O.S., LvZ., S.K., W.B., V.M., J.B., and M.F.Y. conceptualized the study. All authors gave feedback on the manuscript.

## Competing interests

The authors declare no competing interests.

## Methods

### Animals

C57BL/6J (C57BL/6JRj) mice (male, 2.5 months of age) were obtained from Janvier (France). Mice were maintained in a temperature-and humidity-controlled facility on a 12-h reversed light-dark cycle (lights on at 08:15 am) with food and water ad libitum. Mice were housed in groups of 5 per cage and used for experiments when 2.5–4 months old. For each experiment, mice of the same age were used in all experimental groups to rule out confounding effects of age. All tests were conducted during the animals’ active (dark) phase from 12–5 pm. Mice were single housed 24 h before behavioral testing in order to standardize their environment and avoid disturbing cage mates during testing. The animal procedures of these studies were approved by the local veterinary authorities of the Canton Zurich, Switzerland, and carried out in accordance with the guidelines published in the European Communities Council Directive of November 24, 1986 (86/609/EEC).

### Acquisition of mouse data

For mouse behavioral data and annotation, we refer to Sturman et al.^20^. For each day, we randomized the recording chamber of mice used. On days 1-2, we recorded animals 1-8 individually. On day 3, for measuring the effect of interventions on performance, mice were forced-swim-tested in water for 5 minutes immediately before the recording sessions.

### Acquisition of primate data

4 male rhesus macaques were recorded with a 1080p camera within their home-cage. The large indoor room was about 15m^2^. Videos were acquired using a Bosch Autodome IP starlight 7000 HD camera with 1080p resolution at 50 Hz.

### Annotation of segmentation data

To generate training data for segmentation training, we randomly extracted frames of mouse and primate videos using a standard video player. Next, we used the VIA video annotator^39^ to draw outlines around the animals.

### Generation and annotation of primate behavioral videos

For creating the dataset, 3 primate videos of 20-30 minutes were annotated using the VIA video annotator^39^. These videos were generated by previous outputs of SIPEC:SegNet and SIPEC:IdNet. Frames of primates, identified as the same over consecutive frames, were stitched together to create individualized videos. To generate videos of social interactions, we dilated the frames of each primate in each frame and checked if their overlap crossed a threshold, in which case we recalculated the COM of those two masks and center-cropped the frames around them. Labeled behaviors included ‘searching’, ‘object interacting’, ‘social grooming’ and ‘none’ (background class).

### Tracking by segmentation and greedy mask-matching

Based on the outputs of the segmentation masks, we implemented greedy-match-based tracking. For a given frame the bounding box of a given animal is assigned to the bounding box previous frames with the largest spatial overlap, with a decaying factor for temporally distant frames. The resulting overlap can be used as a confidence of SIPEC:SegNet based tracking of the individual. This confidence can be used as a weight when using the resulting track identities to optionally smooth the labels that SIPEC:IdNet.

### Identification labeling with the SIPEC toolbox

As part of SIPEC we release a GUI that allows to label for identification when multiple animals are present (Supp. Fig. 3). To use the GUI, SIPEC:SegNet has to be trained and inference has to be performed on videos to be identity labeled. SIPEC:SegNet results can then be loaded from the GUI and overlaid with the original videos. Each box then marks an instance of the species that is to be labeled in green. For each animal, a number on the keyboard can be defined, which corresponds to the permanent ID of the animal. This keyboard number is then pressed, and the mask-focus jumps to the next mask until all masks in that frame are annotated. Subsequently, the GUI jumps to the next frame in either regular intervals or randomly throughout the video, as predefined by the user. Once a predefined number of masks is reached, results are saved, and the GUI is closed.

### SIPEC top-down workflow

For a given image, if we assume that *N* individuals (with *N* the total number of animals or less) are in the field of view (FOV), the output of SIPEC:SegNet is *N* segmentations or masks of the image. This step is mandatory if the analysis is for multiple animals in a group since subsequent pipeline parts are applied to the individual animals. Based on the masks, the individual animals’ center of masses (COMs) are calculated as a proxy for the animals’ 2D spatial positions. Next, we crop the original image around the COMs of each animal, thus reducing the original frame to *N* COMs and *N* square-masked cutouts of the individuals. This output can then be passed onto other modules.

### SIPEC:SegNet network architecture and training

SIPEC:SegNet was designed by optimizing the Mask R-CNN architecture. We utilized a ResNet101 and feature pyramid network (FPN)^46^ as the basis of a convolutional backbone architecture. These features were fed to the region proposal network (RPN), which applies convolutions onto these feature maps and proposes regions of interest (ROIs). Subsequently, these are passed to a ROIAlign layer, which performs feature pooling, while preserving the pixel-correspondence in the original image. Per level of this pyramidal ROIAlign layer, we assign an ROI feature map from the different layers of the FPN feature maps. Multiple outputs are generated from the FPN, one of which is classifying if an animal is identified. The regressor head of the FPN returns bounding-box regression offsets per ROI. Another fully convolutional layer, followed by a per-pixel sigmoid activation, performs the mask prediction, returning a binary mask for each animal ROI. The network is trained using stochastic gradient descent, minimizing a multi-task loss for each ROI:

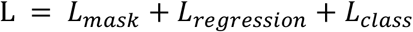

where *L*_*mask*_ is the average binary cross-entropy between predicted and ground truth segmentation mask, applied to each ROI. *L*_*regression*_ is a regression loss function applied to the coordinates of the bounding boxes, modified to be outlier robust as in the original Fast R-CNN paper^47^. *L*_*class*_ is calculated for each of the proposed ROIs (or anchors) as a logarithmic loss of non-animal vs animal. The learning rate was adapted by an animal specific schedule and training was done iteratively, by first training the output layers for some epochs and then incrementally including previous blocks in the training process. SIPEC:SegNet outputs segmentation masks and bounding boxes to create cutouts or masked cutouts of individual animals to be used by one of the downstream modules.

### SIPEC:IdNet network architecture and training

SIPEC:IdNet was based on the DenseNet architecture^28^ for frame-by-frame identification. It consists of 4 dense blocks, which consist of multiple sequences of a batch normalization layer, a ReLU activation, and a convolution. The resulting feature maps are concatenated to the outputs of the following sequences of layers (skip-connections). The resulting blocks are connected through transitions, that are convolutional followed by pooling layers. After the last dense block, we connect an average pooling layer to a Dropout^48^ layer with a dropout rate of 0.5 followed by the softmax classification layer. For the recurrent SIPEC:IdNet, we remove the softmax layer and feed the output of the average pooling layers for each time point into a batch normalization layer^49^ followed by 3 layers of bidirectional gated recurrent units^29,30^ with leaky ReLU activation^50,51^ (alpha=0.3) followed by a Dropout^48^ layer with rate 0.2 followed by the softmax layer. The input for SIPEC:IdNet is the output cutouts of individuals, generated by SIPEC:SegNet (for the single-animal case background-subtracted thresholding and centered-cropping would also work). For the recurrent case, the masks of past or future frames are dilated with a frames per second (FPS) dependent factor that increases with distance in time in order to increase the field of view. We first pre-trained the not-recurrent version of SIPEC:IdNet using Adam^52^ with an lr=0.00025, a batch size of 16 and using a weighted cross-entropy loss. We used a learning rate scheduler in the following form:

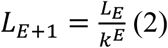

where E stands for epoch, using a k=1.5. Subsequently, we removed the softmax layer and fixed the network’s weights. We then trained the recurrent SIPEC:IdNet again using Adam^52^ and an lr=0.00005, k=1.25 and a batch size of 6.

### SIPEC:BehaveNet network architecture and training

SIPEC:BehaveNet was constructed as a raw-pixel action recognition network. It consists of a feature recognition network that operates on a single frame basis and a network, which integrates these features over time. The feature recognition network (FRN) is based on the Xception^32^ architecture, consisting of an entry, middle, and exit flow. The entry flow initially processes the input with convolution and ReLU blocks. Subsequently, we pass the feature maps through 3 blocks of separable convolution layers, followed by ReLU, separable convolution, and a max-pooling layer. The outputs of these 3 blocks are convolved and concatenated and passed to the middle flow. The middle flow consists of 8 blocks of ReLU layers followed by a separable convolution layer. The Exit receives the feature maps from the middle flow and passes it one more entry-flow-like block, followed by separable convolution and ReLU units. Finally, these features are integrated by a global average pooling layer, followed by a dense layer and passed through the softmax activation. This FRN was first pre-trained on a frame-by-frame basis using an lr=0.00035, gradient clipping norm of 0.5, and batch size=36 using the Adam^52^ optimizer. We reduced the original Xception architecture by the first 17 layers for mouse data to speed up the computation and reduce overfitting. After training the FRN, the outputting dense and softmax layers were removed, and all weights were fixed for further training. The FRN-features were integrated over time by a non-cause Temporal Convolution Network^33^. It is non-causal because, for classification of behavior at time point *t*, it combines features from [*t-n,t+n*] with *n* being the number of timesteps, therefore looking backward in time and forward. In this study, we used an *n* of 10. The FRN features are transformed by multiple TCN blocks of the following form: 1D-Convolution followed by batch normalization, a ReLU activation and spatial dropout. The optimization was performed using Adam^52^ as well with a learning rate of 0.0001 and a gradient clipping norm of 0.5, trained with a batch size of 16.

### Loss adaptation

To overcome the problem of strong data imbalance (most frames are annotated as ‘none’, i.e. no labeled behavior), we used a multi-class adaptation technique Focal loss^53^, commonly used for object detection, and adapt it for action recognition, to discount the contribution of the background class to the overall loss:

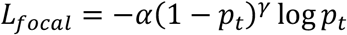

We used a gamma = 3.0 and an alpha = 0.5. For evaluation, we used the commonly used *F1* metric to assess multi-class classification performance while using *Pearson Correlation* to assess temporal correlation.

### SIPEC:PoseNet network architecture and training

Combined with SIPEC:SegNet we can perform top-down pose estimation with SIPEC:PoseNet. That means, instead of the pose estimation network outputting multiple possible outputs for one landmark, corresponding to different animals, we can first segment different animals and then run SIPEC:PoseNet per animal on its cropped frame. In principle, every architecture can now be run on the cropped animal frame, including DLC^2^. The SIPEC:PoseNet architecture is based on an encoder-decoder design^40^. In particular, we used EfficientNet^41^ as a feature detection network for a single frame. Subsequently, these feature maps are deconvolved into heatmaps that regress towards the target location of that landmark. Each deconvolutional layer is followed by a batch normalization layer and a ReLU activation function layer. For processing target images for pose-regression, we convolved pose landmark locations in the image with a 2D Gaussian kernel. Since there were many frames with an incomplete number of labels, we defined a custom cross-entropy-based loss function, which was 0 for non-existing labels.

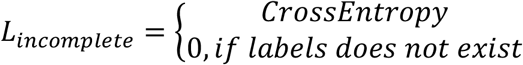

### Combined Model

To test performance effects of doing a pose-estimation-based classification in conjunction with SIPEC:BehaveNet, we pre-trained SIPEC:PoseNet (with classification layer on top) as well as SIPEC:BehavNet individually. Subsequently removed the output layers and fixed the weights of the individual networks and trained a joint output model, which combined inputs of each stream followed by a batch normalization layer, a dense layer (64 units), and a ReLU activation layer. The resulting units were concatenated into a joint tensor followed by a batch normalization layer, a dense layer (32 units), and a ReLU activation layer. This layer was followed by a dense layer with 4 units for the 4 behavioral classes and softmax activation function. This combined model was trained using Adam^52^ with a lr=0.00075. We further offer to use optical flow as an additional input, which has been shown to enhance action recognition performance^54^.

### Implementation and Hardware

For all neural network implementations, we used Tensorflow^55^ and Keras^56^. Computations were done on either NVIDIA RTX 2080 Ti or V100 GPUs.

### 3D location labeling

To annotate the 3D location of a primate, we firstly create a precise model of the physical room (Supp. Fig. 8) using Blender. For a given mask-cutout of a primate, we place an artificial primate at an approximate location in the 3D model. We can then directly read out the 3D position of the primate. 300 samples are annotated, covering the most frequent parts of the primate positions.

### 3D location estimation

To regress the animal positions in 3D, we trained a manifold embedding using Isomap^45^ using the mask size (normalized sum of positively classified pixels), the x and y pixel positions and their pairwise multiplications as features. We used the resulting 6 Isomap features, together with the inverse square root of the mask size, mask size and x-y-position in pixel space to train an ordinary least squares regression model to predict the 3D position of the animal.

### Metrics used

Abbreviations used: Pearson – Pearson Correlation, RMSE – Root mean squared error, IoU – intersection over union, mAP – mean average precision, dice – dice coefficient.

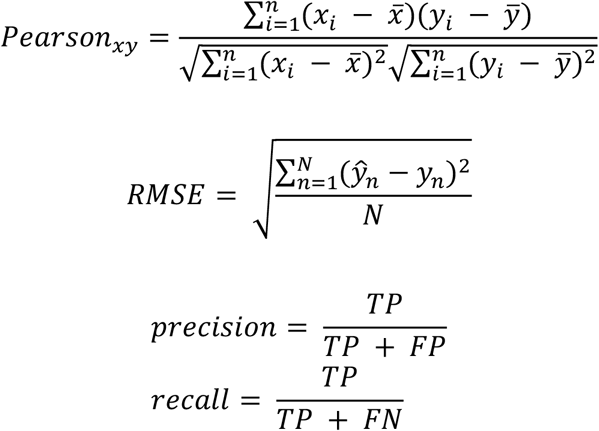

Where TP denotes True Positives, FP False Positives, TN True Negatives, and FN False Negatives.

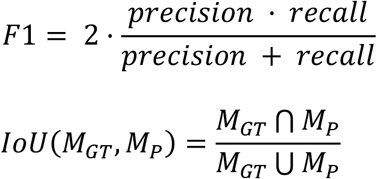

Where *M*_*GT*_ denotes the ground truth mask and *M*_*P*_ the predicted one. We now calculate the mAP for detections with an IoU > 0.5 as follows:

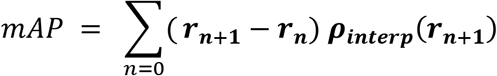

With

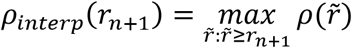

Where *ρ*(*r*) denotes precision measure at a given recall value.

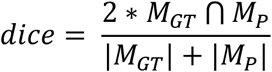

## Supplementary

**Supplementary Fig. 1.**
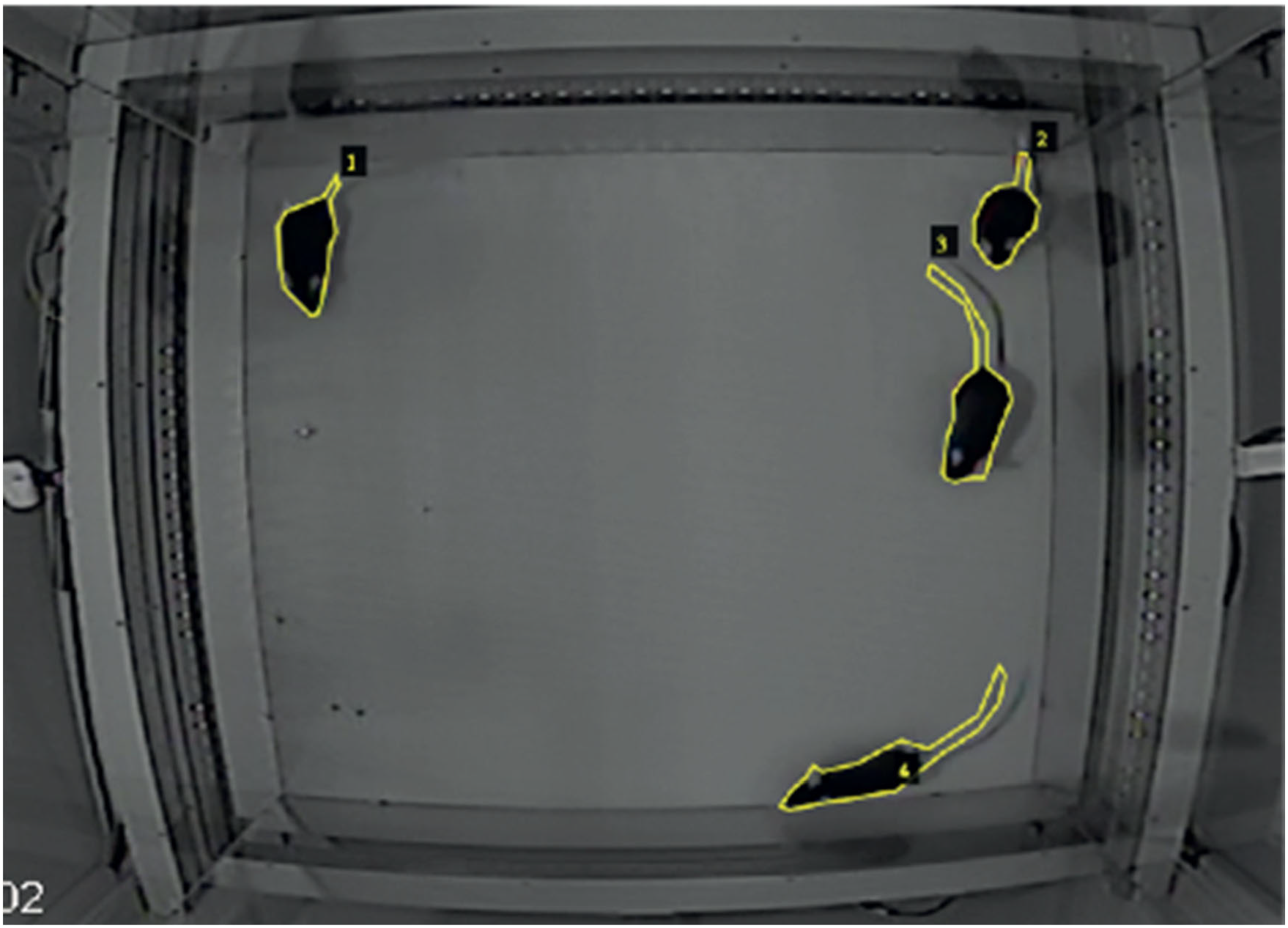
Segmentation annotation illustration. An exemplary frame of mice in OFT with manually annotated outlines.

**Supplementary Fig. 2.**
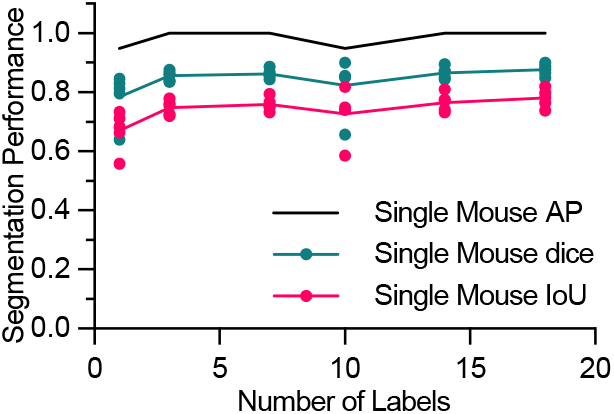
Mouse single segmentation. For mice, SIPEC:SegNet performance in mAP, dice and IoU for single mouse as a function of the number of labels. The lines indicate the means for 5-fold CV while circles, squares, triangles indicate the mAP, dice, and IoU, respectively, for individual folds. All data is represented by mean, showing all points.

**Supplementary Fig. 3.**
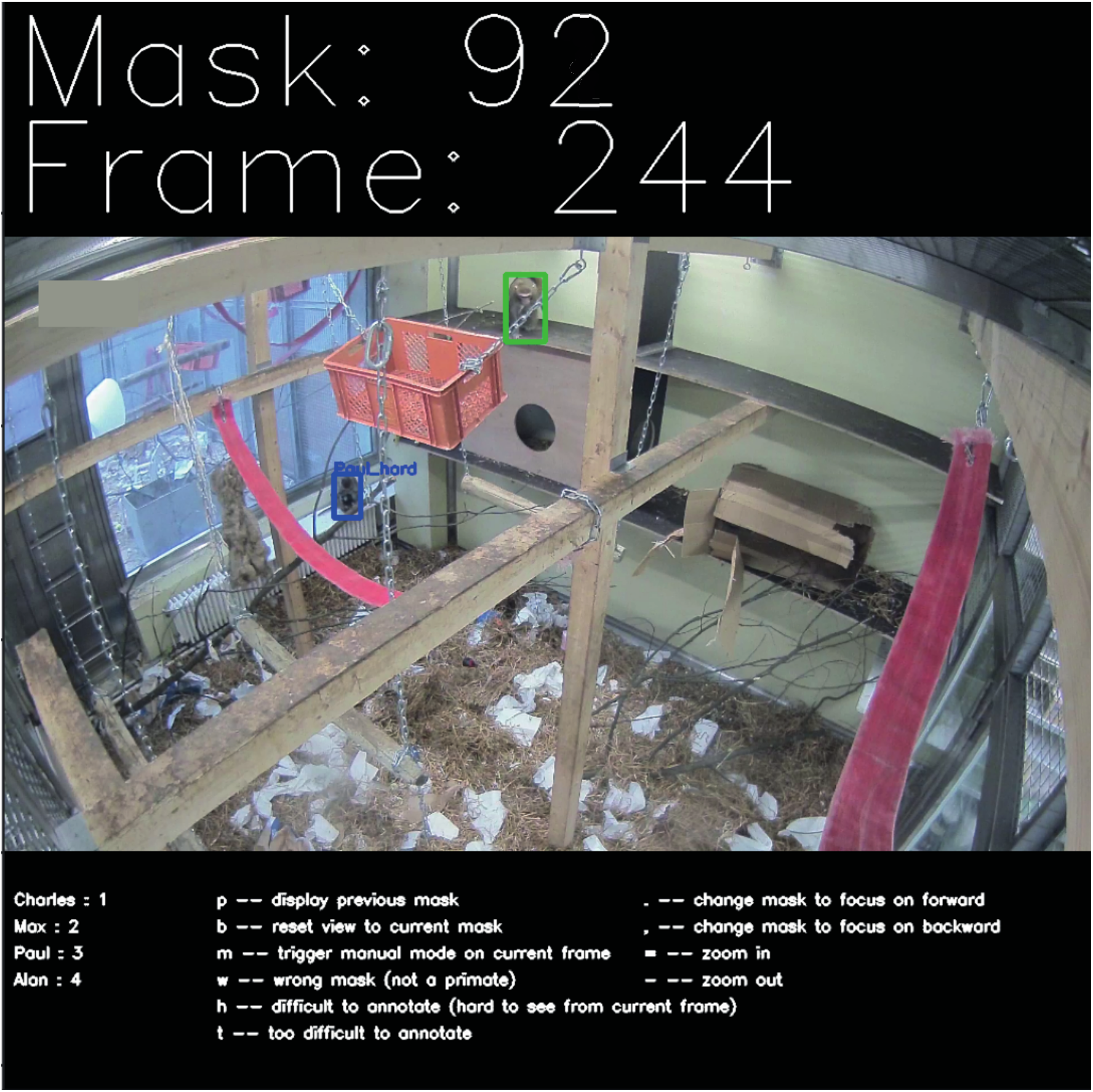
Identification Graphical User Interface. Mask-box results from SIPEC:SegNet is overlaid over frames in blue and can be labeled one by one. The current box to be labeled is in green. A simple keyboard input scheme is provided within the GUI. Names of individuals and the number of masks to be labeled can be set by the user.

**Supplementary Fig. 4.**
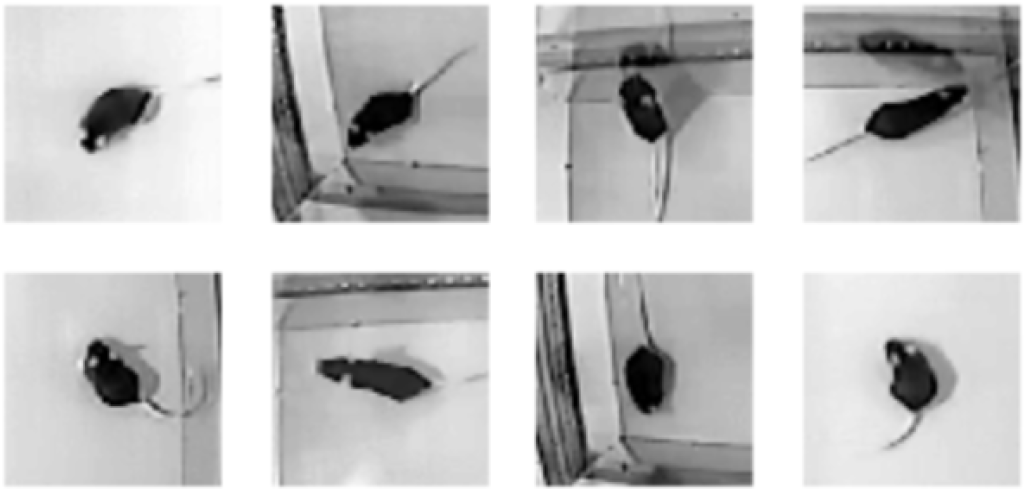
Example frames of the 8 distinct mice.

**Supplementary Fig. 5.**
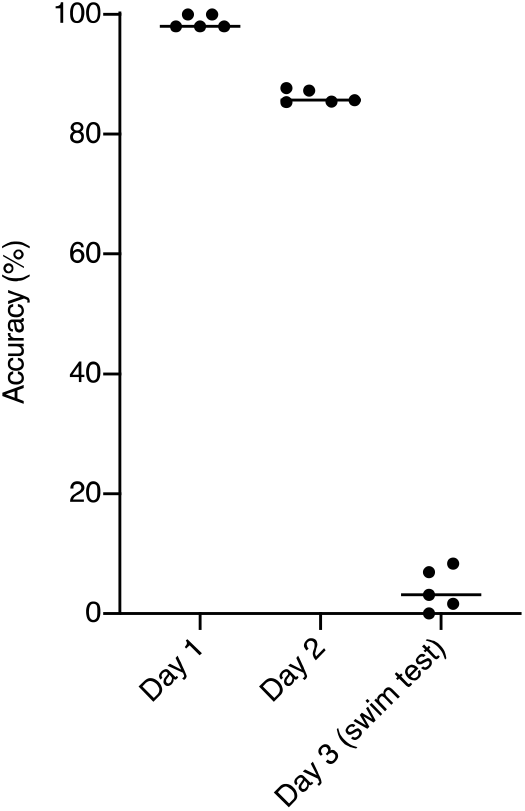
Identification performance of mice across days and interventions. Identification accuracy across days for models trained on day 1. While the performance for the day the model is trained on is very high it drops when tested on day 2, but is still significantly above chance level. When tested on day 3, after a forced swim test intervention, the performance drops significantly. All data is represented by mean, showing all points.

**Supplementary Fig. 6.**
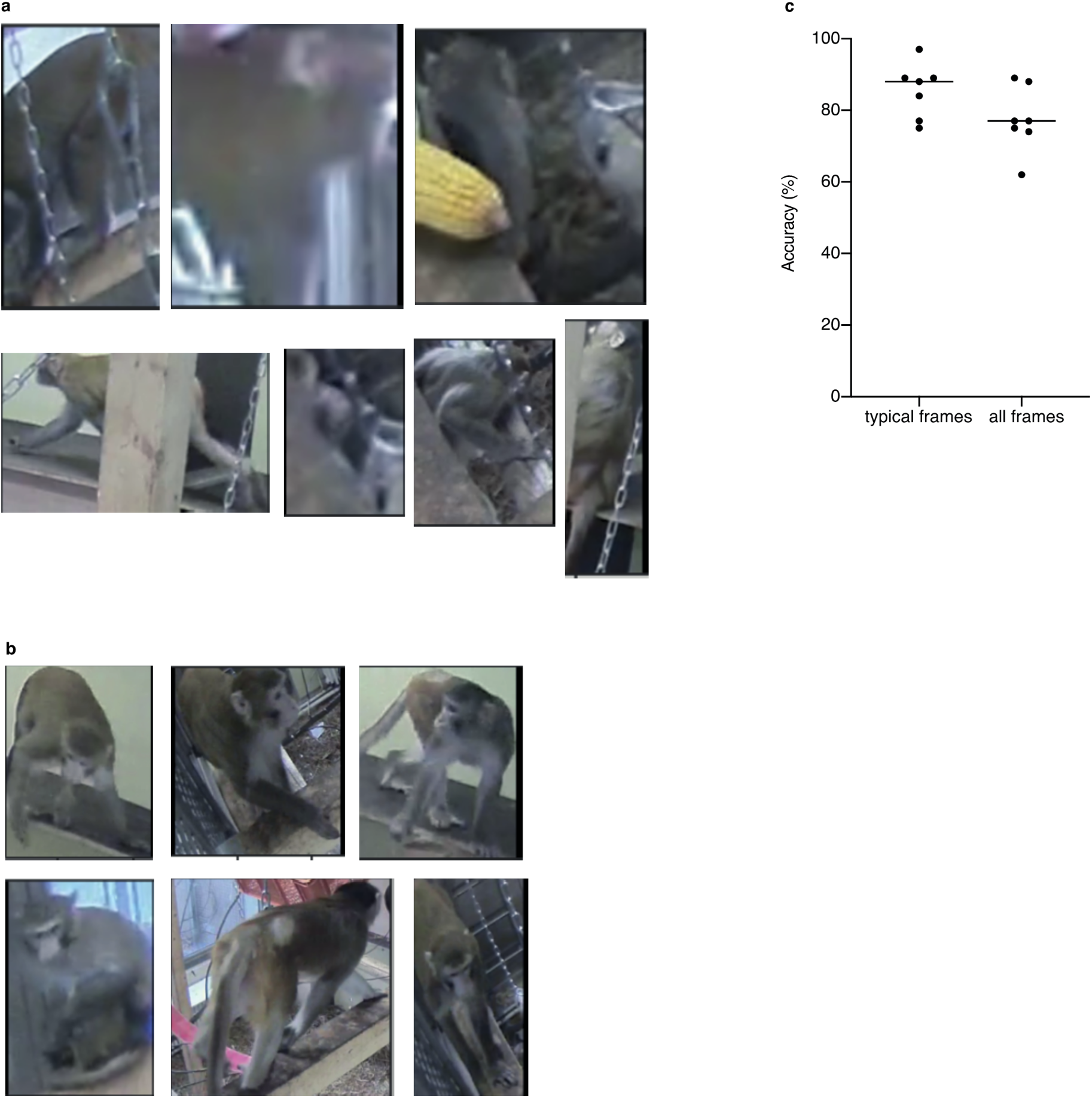
Identification of typical vs difficult frames. a) Very difficult exemplary frames, which are also beyond human single-frame recognition, are excluded for the ‘typical’ frame evaluation. b) Exemplary frames used for the ‘typical’ frame analysis. c) Identification performance is significantly higher on ‘typical’ frames than on all frames. All data is represented by mean, showing all points.

**Supplementary Fig. 7.**
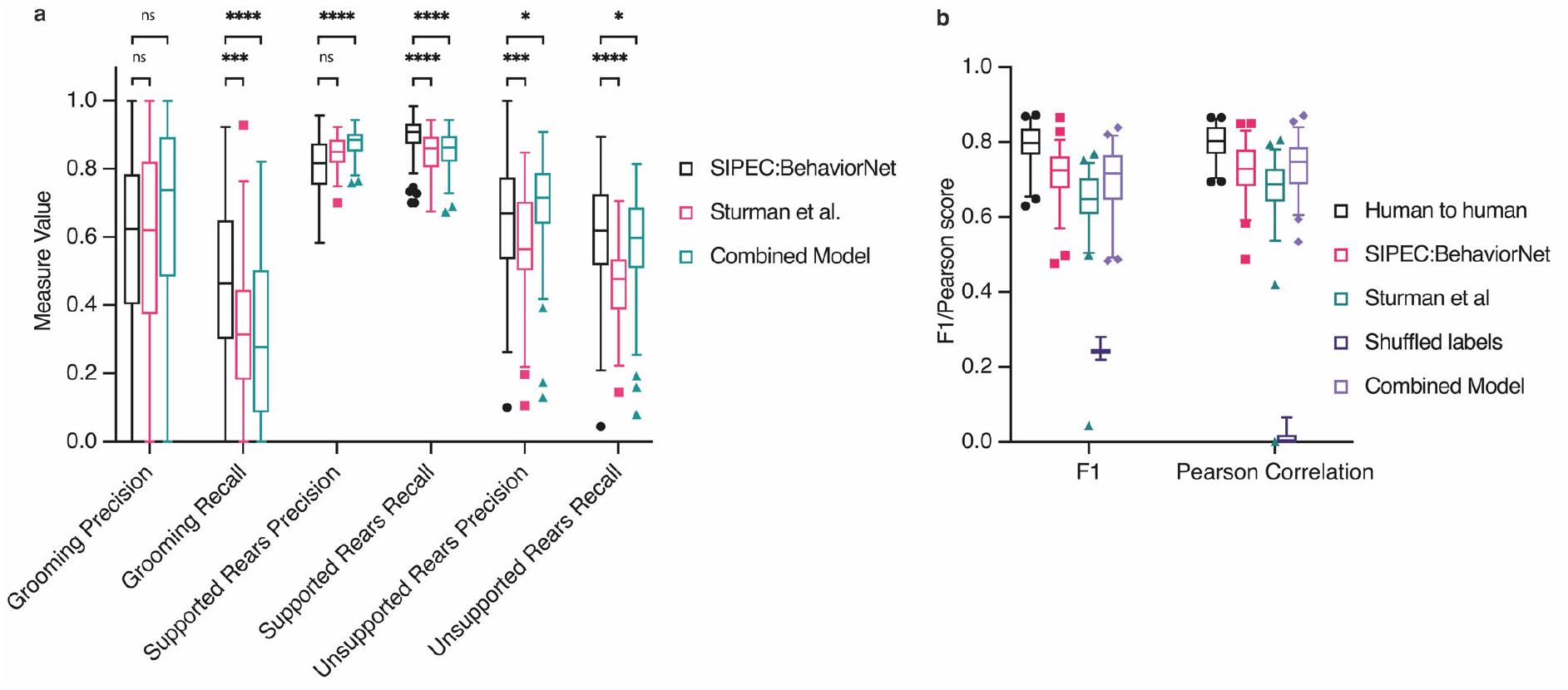
Additional behavioral evaluation. a) Overall increased F1 score is caused by an increased recall in case of grooming events and precision for unsupported rearing events. b) Comparison of F1 values as well as Pearson Correlation of SIPEC:BehaveNet to human-to-human performance as well as combined model. Using pose estimates in conjunction with raw-pixel classification increases precision in comparison with solely raw-pixel classification while suffering from a decrease in recall. All data is represented by a Tukey box-and-whisker plot, showing all points.

**Supplementary Fig. 8.**
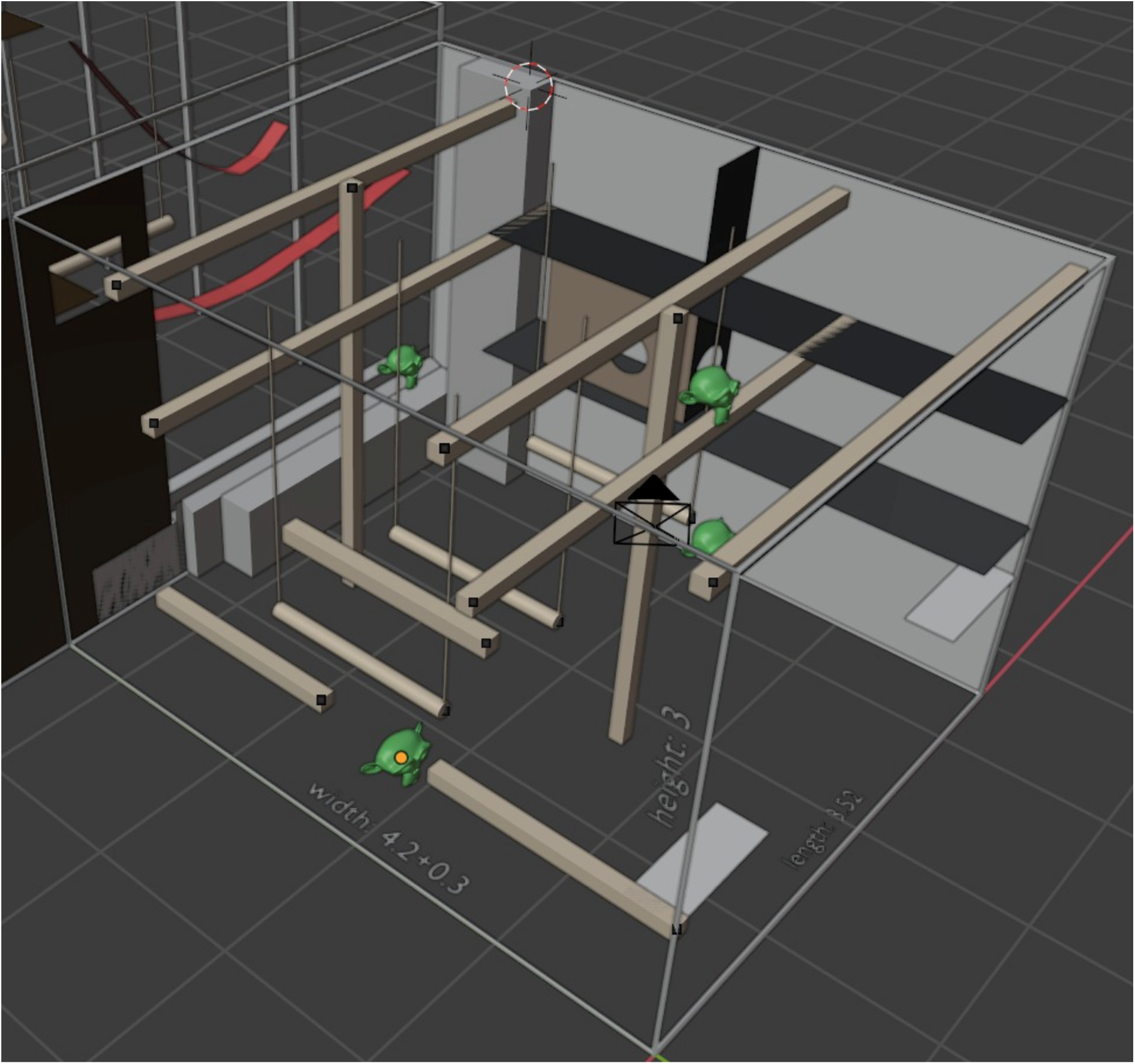
3D model used for annotation of primate 3D-location data.

**Supplementary Fig. 9.**
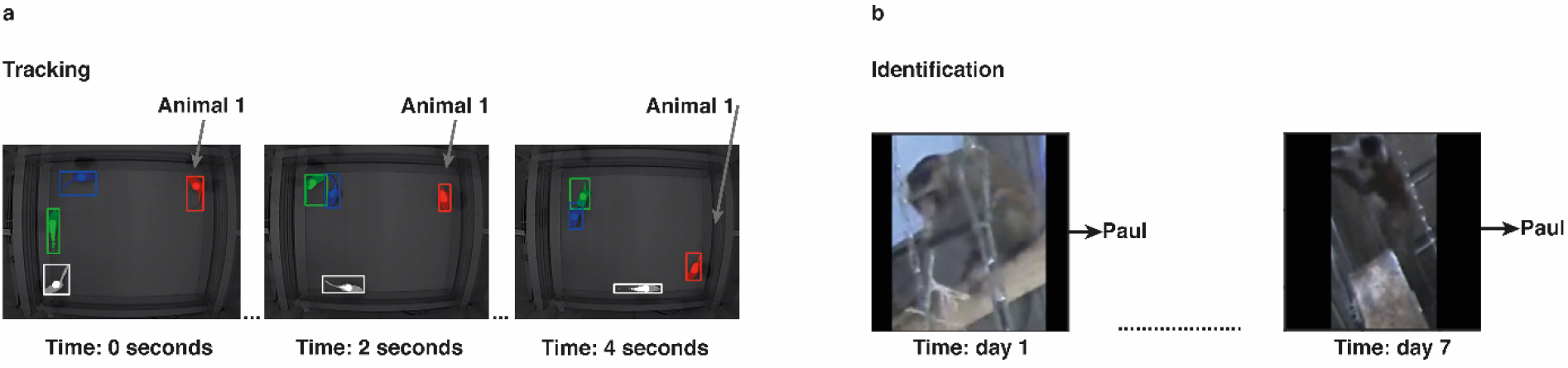
Comparison tracking and identification. a) **Tracking** describes the process of following each individual animal in a group of animals within one session in a given field of view. b) **Identification** describes the ability to identify an individual from a single frame or a few consecutive frames across multiple sessions that could be apart hours, days, or months. This entails difficulties such as varying lighting conditions, occlusions, changes in the appearance of animals over time.

**Supplementary Fig. 10.**
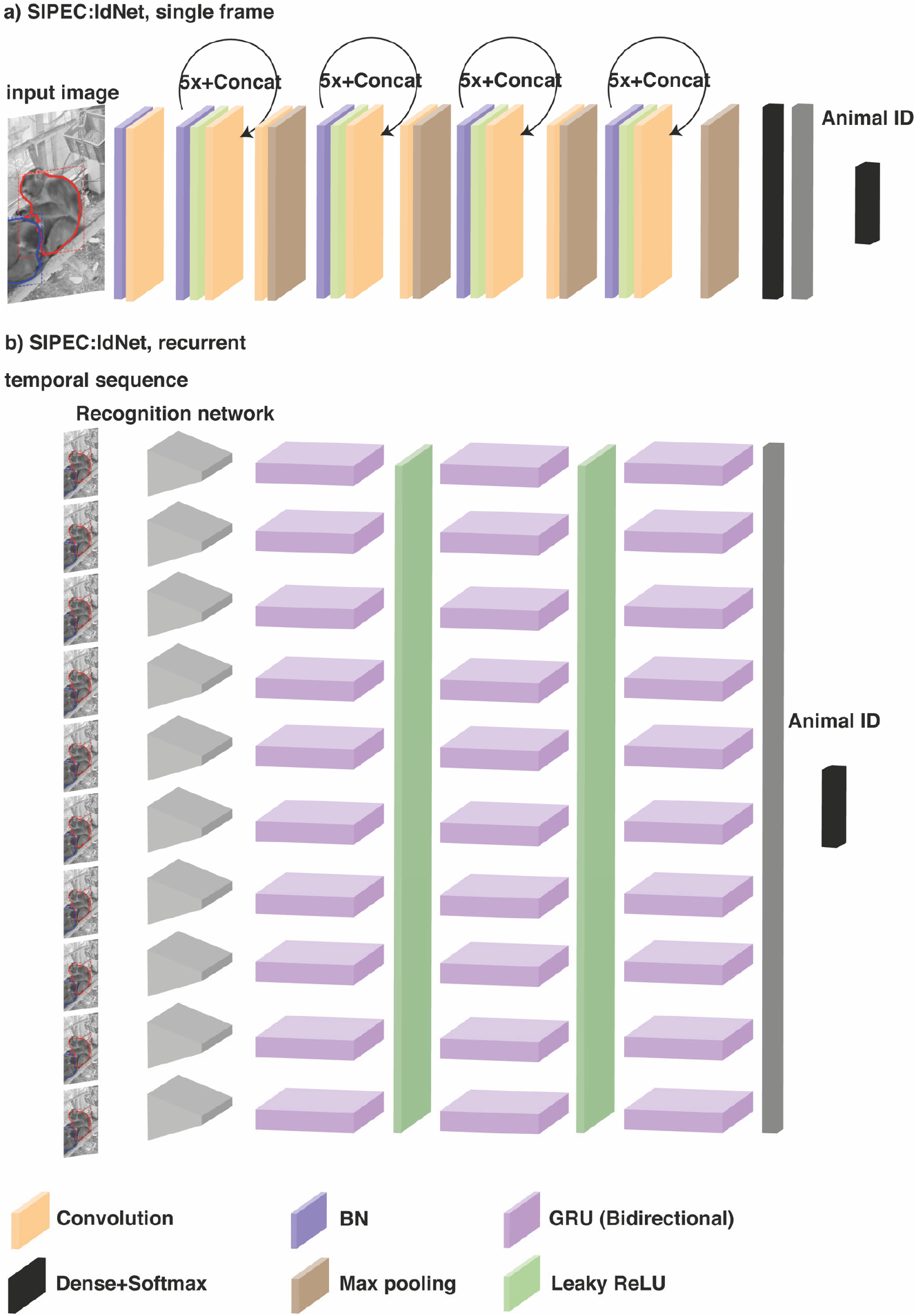
SIPEC IdNet Architecture.

**Supplementary Fig. 11.**
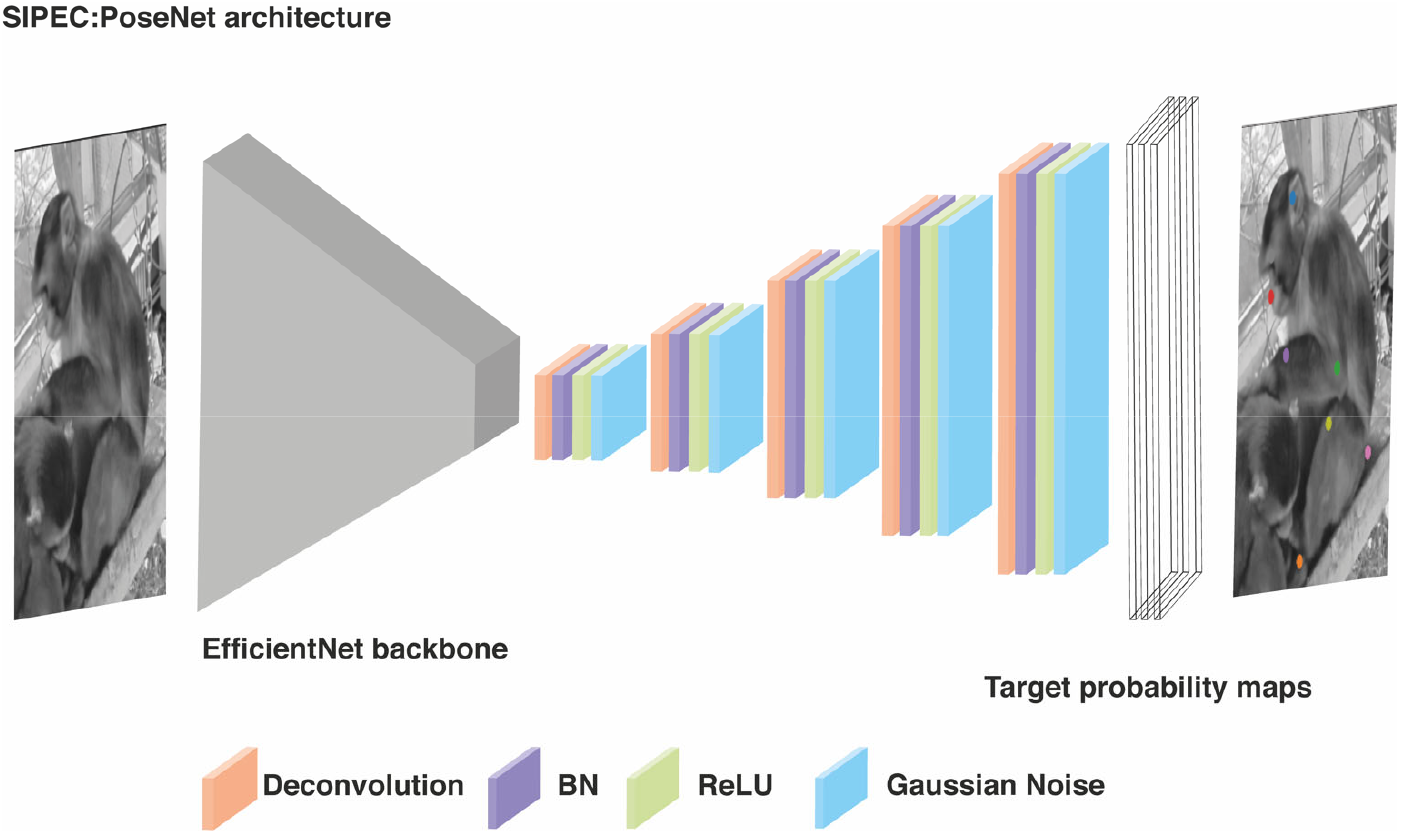
SIPEC PoseNet Architecture.

**Supplementary Fig. 12.**
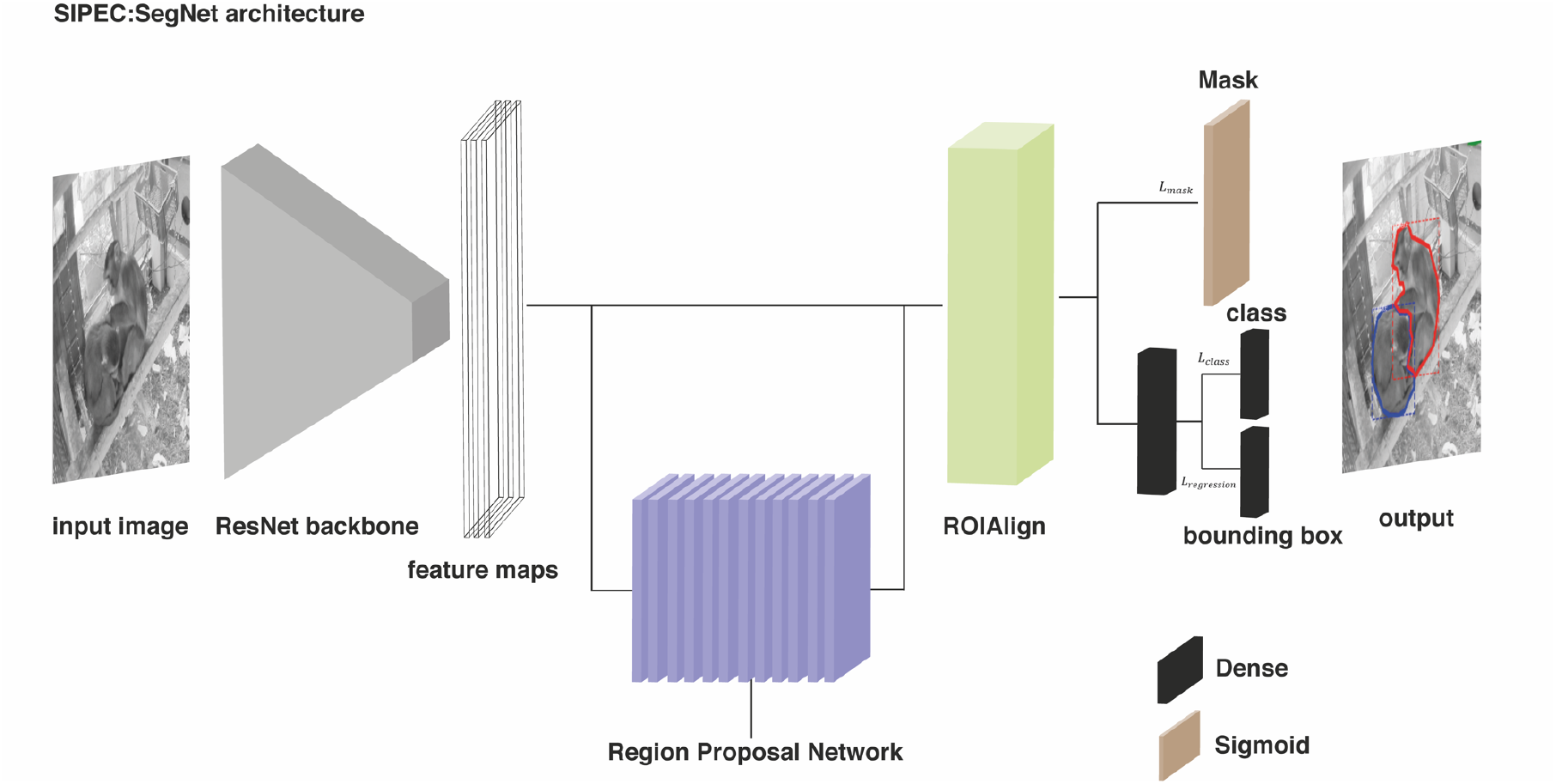
SIPEC SegNet Architecture.

**Supplementary Fig. 13.**
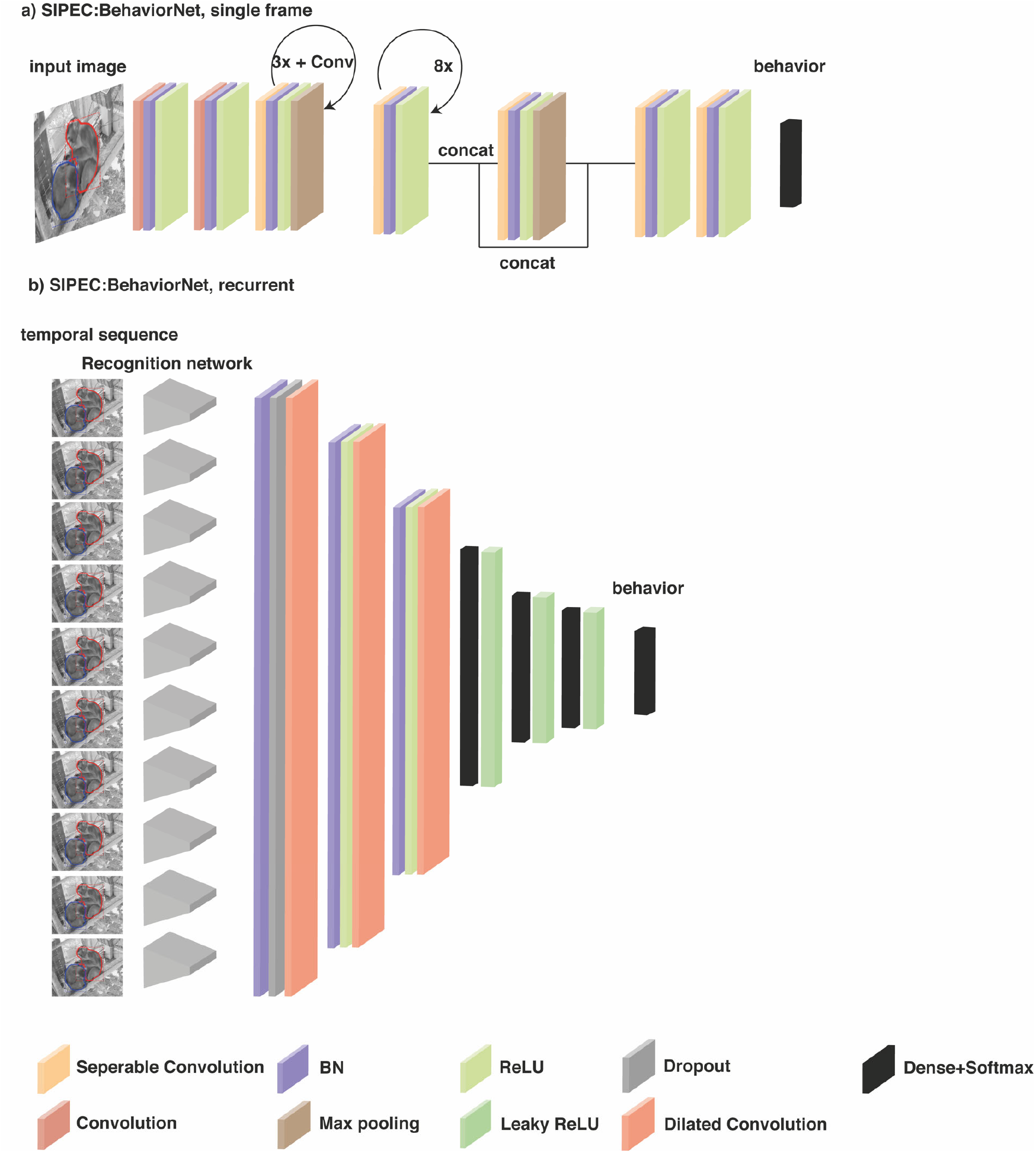
SIPEC BehaviorNet Architecture.

**Supplementary Fig. 14.**
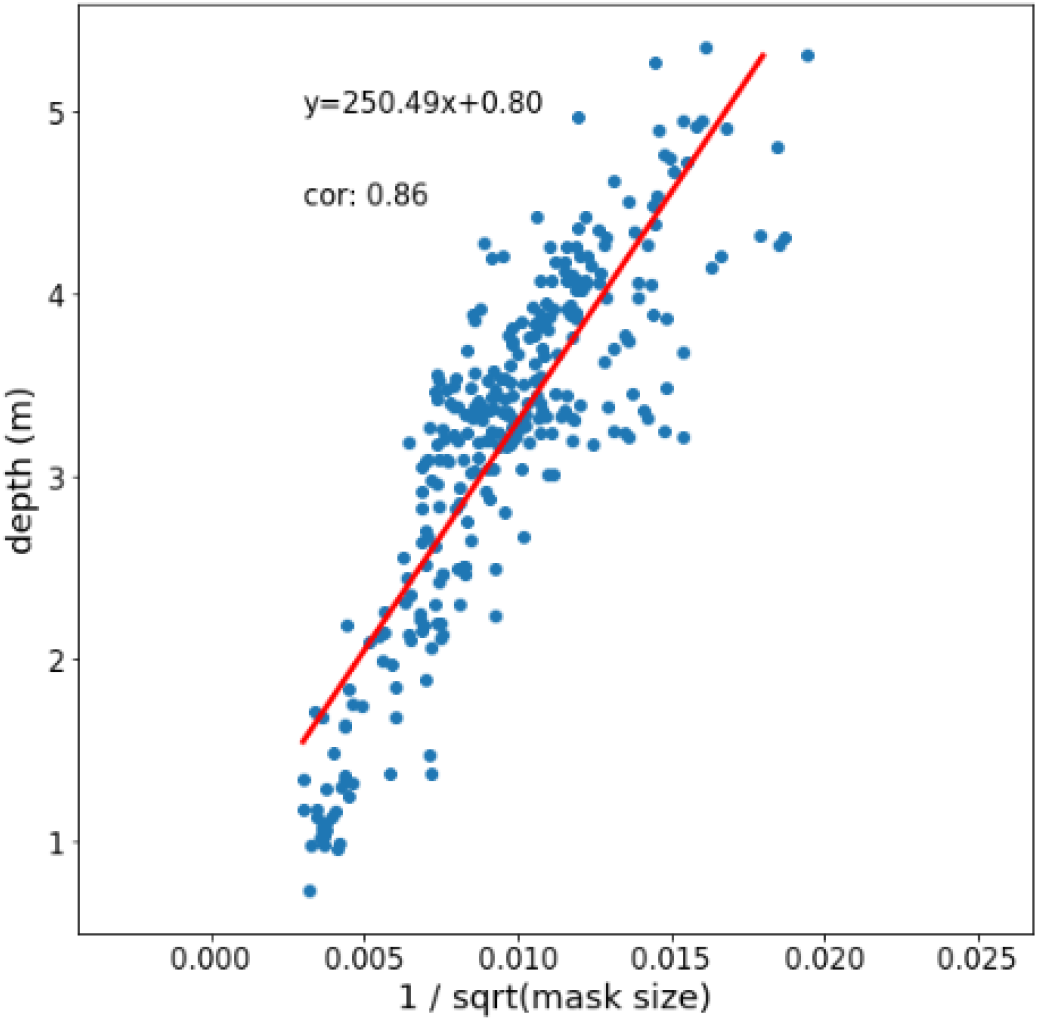
3D depth estimates based on mask size. The inverse of the square root of the mask size (based on SIPEC:SegNet output) highly correlates with the depth of the

**Supplementary Fig. 15.**
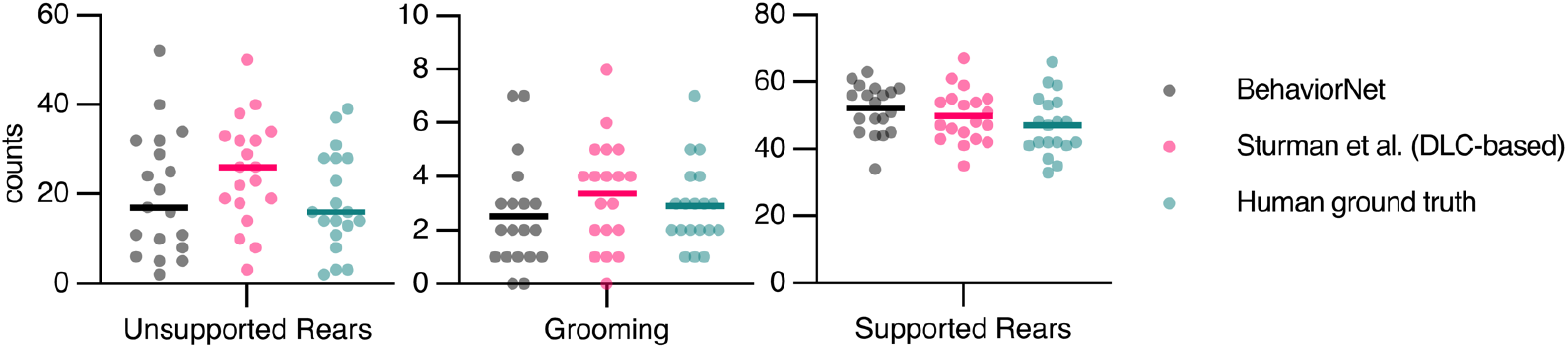
Comparison of counts of behaviors between SIPEC:BehaviorNet, pose estimation based approach and human raters. Unsupported and supported rears and grooming events were counted per video for n=20 different mice videos. Behaviors were integrated over multiple frames, as described in Sturman et al.^14^. Behavioral counts of 3 different human expert annotators were averaged (in legend as ‘human ground truth’). No significant differences were found for comparing the number of behaviors between SIPEC:BehaviorNet and human annotators or Sturman et al.^14^ and human annotators (Tukey’s multiple comparison test). All data is represented by mean, showing all points.

**Supplementary Tab. 1.** Training and inference times. All measures are done with an NVIDIA RTX 2080 Ti and represent average values.

| Species | Network | Training (seconds/epoch) | Epochs | Total training (min) | Inference (seconds/frame) |
| --- | --- | --- | --- | --- | --- |
| Primate | SegNet | 133 | 100 | 222 | 0.4 |
| Primate | IdNet(single frame) | 134 | 10 | 22 | 0.35 |
| Primate | IdNet(recurrent) | 60 | 20 | 20 | 0.6 |
| Primate | PoseNet | 34 | 900 | 510 | 0.07 |
| Primate | BehaveNet(single frame) | 15 | 50 | 5 | 0.08 |
| Primate | BehaveNet(recurrent) | 190 | 10 | 31 | 0.6 |
| Mouse | SegNet | 40 | 100 | 67 | 0.14 |
| Mouse(@1000 frames) | IdNet(single frame) | 432 | 10 | 72 | 0.1 |
| Mouse(@1000 frames) | IdNet(recurrent) | 502 | 10 | 83 | 0.19 |
| Mouse | PoseNet | 20 | 2450 | 817 | 0.03 |
| Mouse | BehaveNet(single frame) | 547 | 10 | 91 | 0.05 |
| Mouse | BehaveNet(recurrent) | 1163 | 10 | 194 | 0.2 |

**Supplementary Video 1** | **Illustration of SIPEC:SegNet and SIPEC:IdNet in primate homecage environment**.

Short exemplary video of behaving primates in their homecage environment. SIPEC:SegNet is used to mask different primates and SIPEC:IdNetis used to identify them. During obstructions, the identity of a primate can alter but SIPEC:IdNet quickly recovers the correct identity over the next frames, as it becomes more visible and therefore better identifiable.

**Supplementary Video 2** | **Comparison of SIPEC and idtracker.ai for mice**.

Comparison for tracking 4 mice by idtracker.ai (Left) and by SIPEC(Right). We used publicly available data from idtracker.ai (https://drive.google.com/drive/folders/1Vua7zd6VuH6jc-NAd1U5iey4wU5bNrm4) as well as idtracker.ai’s publicly available inference results (https://www.youtube.com/watch?v=ANsThSPgBFM) for a tracking comparison. **Left video**: The tracking of idtracker.ai exhibits prolonged label switching errors where the label of two or more animals gets swapped for some time. **Right Video**: Tracking is performed by SIPEC:SegNet in conjunction with greedy-mask matching to track the identities of animals. In this example video, SIPEC is more robust to these kinds of errors than idtracker.ai. (see also Supp. Video 4).

**Supplementary Video 3** | **Tracking of 4 mice by SIPEC in an open-field test**.

The masks generated by SIPEC:SegNet in conjunction with greedy-mask matching are used to robustly track identities of four mice in an open-field test (see Methods).

**Supplementary Video 4** | **SIPEC tracking over 52-minute video**.

We used publicly available data from idtracker.ai (https://drive.google.com/drive/folders/1Vua7zd6VuH6jc-NAd1U5iey4wU5bNrm4) and tracked 4 mice. The masks generated by SIPEC:SegNet in conjunction with greedy-mask matching are used to robustly track identities of four mice in an open-field test (see Methods).

## Notes

https://github.com/SIPEC-Animal-Data-Analysis/SIPEC

### Competing Interest Statement

The authors have declared no competing interest.

### Summary of Updates

Updated manuscript and Supplementary; additional Supplementary Videos

